# Mutation bias influences the emergence and effects of antibiotic resistance

**DOI:** 10.64898/2026.08.18.745421

**Authors:** Adrita Chakraborty, Deepa Agashe

## Abstract

Evolution of antibiotic resistance is a major global public health problem. Rapid emergence of antibiotic resistance is often linked to hypermutator bacteria with defective DNA repair, leading to high mutation rates that are broadly advantageous. However, depending on which DNA repair pathway is dysfunctional, mutators may sample only specific types of mutations at a higher rate. Thus, their mutation spectrum can be biased towards specific mutation types, influencing the identity of resistance mutations. Under strong antibiotic selection, an overall high mutation rate should generally shorten the time to sample a resistance mutation and increase the probability of resistance. However, recent work suggests that the mutation rate for specific types of mutations in target genes that drive high resistance is more important than the overall mutation rate. To systematically test this prediction, we exposed *Escherichia coli* mutators with varying mutation rates and spectra to antibiotics targeting different cellular functions. For each strain, we determined the highest antibiotic concentration at which resistance could emerge overnight, quantifying both the magnitude and probability of resistance. High level antibiotic resistance was generally better predicted by specific rather than overall mutation rate, and resistance mutations matched the mutation spectrum of the respective mutator. Despite the varying magnitude of resistance, at the highest antibiotic concentration survived by each strain, the respective resistance mutations were generally costly in the absence of antibiotic. Given that mutators often arise in laboratory, natural, and clinical settings under antibiotic selection, we suggest that their mutation spectra deserve more attention.

## INTRODUCTION

Soon after the introduction of antibiotics in the mid-20^th^ century, antimicrobial resistance (AMR) emerged as a global public health crisis, threatening the effective prevention and treatment of bacterial infections worldwide. In 2019, approximately 4.95 million deaths were attributed to AMR, and this figure is estimated to reach 10 million by 2050 (Murray et al., 2022). To combat AMR, it is necessary to understand the mechanisms, and the ecological and evolutionary factors that increase the likelihood of resistance emergence. Resistance mutations typically alter the drug binding sites of target genes or proteins, upregulate efflux pumps to decrease intracellular concentration of antibiotics, or rewire metabolic and regulatory pathways to offset the drug’s inhibitory or bactericidal effects (Woodford and Ellington, 2007), and therefore spread in the population (MacLean et al., 2010). Such selective sweeps of resistance mutations are indeed observed in patients treated with antibiotics for bacterial infections (Shepherd et al., 2024). The probability of acquiring beneficial resistance mutations is directly correlated with the overall mutation rate, and hence, hypermutator bacterial strains arise frequently in laboratory evolution experiments with antibiotic selection and are a particular concern in clinical settings (LeClerc et al., 1996; Sniegowski et al., 1997; Oliver et al., 2000; Komp Lindgren et al., 2003; Mehta et al., 2019; Gifford et al., 2023). For instance, mutation rate is strongly correlated with clinical resistance against fluoroquinolones via multiple target genes in *Escherichia coli* (Komp Lindgren et al., 2003), with some hypermutators showing up to a 1000-fold higher emergence of resistance against very high antibiotic doses, compared to non-hypermutators (Miller et al., 2002). Increasing the mutation rate also increases the rate of evolutionary adaptation to different antibiotics (Shibai et al., 2025). Additionally, hypermutators are more likely to show multi-drug resistance, because when a mutator allele rises to fixation by hitchhiking with a beneficial resistance mutation, it causes rapid emergence of other resistance mutations (Gifford et al., 2023). However, the benefit of hypermutation is inconsistent across different hypermutators, antibiotics, and doses, indicating a role for additional factors.

Hypermutator bacteria usually have high mutation rates because their DNA damage repair pathways are impaired (Jolivet-Gougeon et al., 2011; Foster et al., 2015; Hall et al., 2025). However, depending on which DNA damage repair gene is affected, the occurrence of only specific types of mutations — that would otherwise be repaired by the deficient gene — is increased. Thus, hypermutators often have biased mutation spectra. For instance, in *E. coli,* the MutT protein scavenges 8-oxo-dGTP, reducing the likelihood of oxidative DNA damage. As a result, deletion of *mutT* leads to a 100-fold increase in the rate of TA→GC transversions (Foster et al., 2015). Thus, if TA→GC mutations are beneficial — e.g., under high doses of Streptomycin (Strep) where high level resistance often involves TA→GC mutations in the target gene *rpsL* — a *ΔmutT* strain should develop resistance more rapidly, compared to mutator strains with a different mutation bias. Indeed, two mutator strains with distinct mutation spectra each had a higher probability of developing resistance to two different antibiotics: *ΔmutY* had a higher rate of resistance with faster growth in Rifamycin (Rif) despite having a lower mutation rate, whereas in Strep, many more resistant clones were found in *ΔmutT*, and they also grew faster (Couce et al., 2013). These effects of mutation bias hold even in large populations where the effect of mutation supply is thought to be weaker (Barber et al., 2025). More generally, strong mutation biases in mutators can dramatically alter the genetic basis of resistance to antibiotics, as observed for *ΔmutH* and *ΔmutT* strains evolving under Cefotaxime (Cef) selection (Couce et al., 2015). An analysis of mutation spectra and adaptive mutations in three different species of bacteria found that the mutation bias of an organism influences which kind of adaptive mutations arise (Cano et al., 2022), reinforcing the idea that the arrival bias of mutations is a major predictor of adaptation (Stoltzfus et al., 2017). Thus, hypermutators may accelerate the emergence of antibiotic resistance not only by increasing the overall supply of mutations, but also by biasing the types of mutations that are frequently sampled under selection. However, because most prior studies focused on a relatively small number of mutators and antibiotics, broad evidence and quantification of the distinct roles of mutation rate vs. specific mutation bias in different antibiotics remains elusive.

To address this gap, we exposed six *E. coli* mutator strains with varying mutation rates (WT (low), intermediate (10X WT) and high (100x WT)) and spectra to six different antibiotics targeting different cellular functions. We quantified their survival in increasing concentrations of antibiotic, rate of emergence of resistance in overnight cultures, and identified newly acquired resistance mutations by sequencing (Fig. 1A-B). We hypothesized that under strong antibiotic selection, high mutation rate should generally facilitate adaptation by reducing the time to sample a beneficial resistance mutation (Fig.1C). Further, strains whose mutation spectra align with known high-benefit resistance mutations in target genes (obtained from Comprehensive Antibiotic Resistance Database (CARD) (Alcock et al., 2023); MicroBIGG-E (Feldgarden et al., 2021); Pelchovich et al., 2013; Jin and Gross, 1988; van der Putten et al., 2019; Froelich et al., 2006; Couce et al., 2015) should be more likely to evolve resistance to very high antibiotic concentrations, by sampling those beneficial resistance mutations more rapidly (Fig. 1D). We find that resistance does not always arise faster in strains with the highest mutation rates. Instead, specific mutation bias (high rate of specific types of mutations) is a better predictor of rapid emergence of antibiotic resistance. Our findings highlight the importance of considering mutation spectra of mutator alleles while studying the evolutionary consequences of antibiotic resistance emergence.

**Fig. 1:**
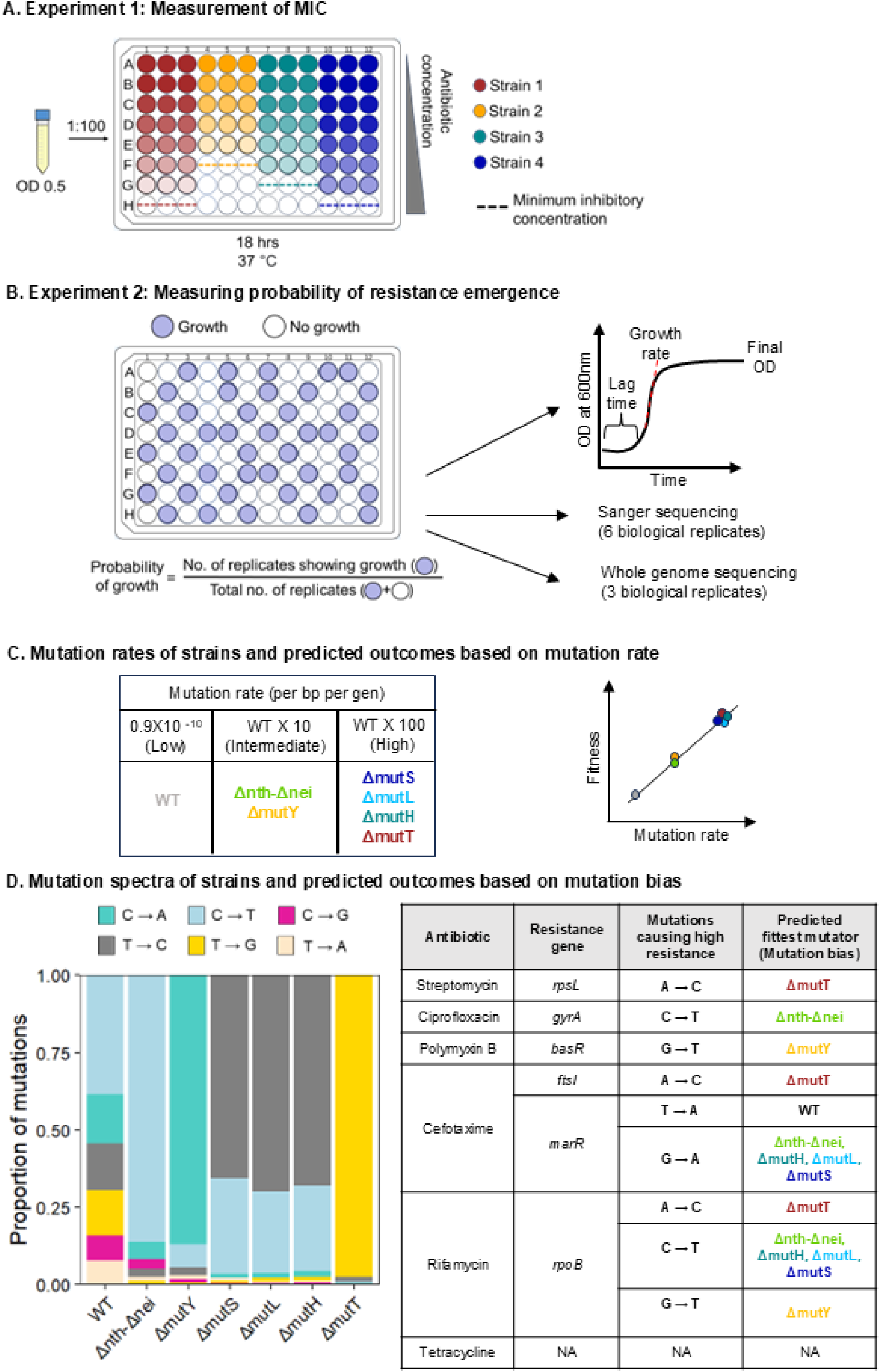
Experimental design and expectations. **(A)** Illustration of the experimental design to measure minimum inhibitory concentration (MIC), probability of emergence of resistance, and growth parameters (lag time, final OD and growth rate); and sequencing. We determined MIC using broth microdilution assays across a gradient of antibiotic concentrations. The MIC for each strain–antibiotic combination was defined as the lowest concentration that prevented visible growth after 18 h (Experiment 1). **(B)** At the highest sub-MIC of each strain-antibiotic combination, we quantified the probability of resistance emergence using a large number of replicate populations cultures in antibiotic-supplemented medium, monitored over a single growth cycle (Experiment 2). Growth/no-growth outcomes (Cutoff for growth = OD_600_ above 0.02) from these OD measurements were used to estimate the probability of resistance emergence. We used growth curves so obtained to estimate lag time, growth rate, and final OD. We randomly selected a few replicate populations for Sanger sequencing (n=6) and whole genome sequencing (n=3). **(C)** Mutation rate of all strains used in this study, and their predicted fitness in all tested antibiotics based on mutation rate. **(D)** Mutation spectra of nucleotide substitutions of all strains as reported in Sane et al., 2025, and predicted fitness in each tested antibiotic based on the specific mutation bias of each strain. For mutation spectra including insertions and deletions look at Fig. S11. For each antibiotic, we used prior reports of known target genes involved in resistance and the specific nucleotide substitutions that confer high-level resistance, to predict which mutator should have the greatest probability of sampling beneficial resistance mutations (Comprehensive Antibiotic Resistance Database (CARD) (Alcock et al., 2023); MicroBIGG-E (Feldgarden et al., 2021); Pelchovich et al., 2013; Jin and Gross, 1988; van der Putten et al., 2019; Froelich et al., 2006; Couce et al., 2015)).

## METHODS

### Bacterial strains

We used wild type (WT) and mutator strains (*ΔmutT, ΔmutH, ΔmutL, ΔmutS, Δnth-Δnei, ΔmutY*) generated as described previously (Sane et al., 2025). Briefly, we obtained the wild-type strain of *E. coli* MG1655 and each mutator strain (in the BW25113 strain background) from the Keio collection of the Coli Genetic Stock Centre (CGSC, Yale University). We moved the knockout locus with a kanamycin resistance cassette from the BW25113 background into the MG1655 background. We removed the kanamycin resistance cassette, leaving an inframe deletion of the desired gene. We PCR-sequenced the knockout locus to confirm complete removal of the kanamycin cassette and successful generation of the mutator strains. We made stocks of all strains, preserved at –80°C (1:1 ratio of overnight grown bacterial culture in Luria Broth (LB) and 60% glycerol).

### Experiment 1: Assay to determine the Minimum Inhibitory Concentration (MIC)

We used six different antibiotics: streptomycin (Strep), polymyxin B (Pol B), ciprofloxacin (Cip), cefotaxime (Cef), rifamycin (Rif) and tetracycline (Tet) (all from Sigma, except Tet from Tokyo Chemicals Limited). We determined the MIC for each antibiotic and strain using the standard broth microdilution assay (Wiegand et al., 2008) in LB (Difco). We inoculated 1 µL of thawed cryostocks in 100 µL of LB in 96 well microplates (Genetix), reviving cultures with orbital shaking at 200 rpm for 12 hours at 37 °C. For the MIC assay, we wanted to ensure that we inoculated similar numbers of cells in each well (2 to 8 x10^5^ per µL) as per Clinical & Laboratory Standards Institute (CLSI) guidelines. We counted colony forming units (CFU/ml) for several batches of revived cultures to confirm that this could be reliably achieved by diluting cultures to a blank-corrected optical density (OD_600_) of 0.5 (Biotek Epoch 2 reader) (Fig. S1). We added 1 µL of the diluted revived culture to antibiotic-supplemented LB, serially increasing the antibiotic concentration (n=3 technical replicates per concentration). For each antibiotic, we covered a wide range of concentrations, from 0 to 2X reported MIC for *E. coli*, with 2-fold step increases in concentration. Based on initial results, we either added more steps (i.e., higher concentrations) or tested smaller concentration steps (<2 fold increases) to try and distinguish between strains. To maintain constant nutrient availability, we always added 1 µL culture and 1 µL antibiotic stock solution to 98 µL LB, using antibiotic stock solutions of different concentrations as required (see Fig. S2 for growth curves and antibiotic concentrations used for MIC measurements). We incubated each microplate for 18 hours at 37 °C with orbital shaking at 425 cpm in a Biotek plate reader, measuring OD_600_ every 15 mins. Each plate included un-inoculated control wells to verify media sterility. We defined the MIC as the antibiotic concentration at which none of the 3 replicates of a strain showed any growth in 18 hours.

To confirm that active antibiotic was present at the end of the 18 hour growth cycle, we measured growth in spent media (Fig. S3). We cultured each strain for 18 hours in LB containing each antibiotic at the highest sub-MIC concentration determined from the MIC assay. We collected spent media by filtering cultures using a sterile 0.22 µm bacterial filter, using spent media of cultures grown in LB without antibiotic as a control, and an additional negative control with uninoculated filtered spent media. We inoculated the spent media with fresh cultures of the corresponding strains and monitored growth for a second growth cycle, as above.

### Experiment 2: Quantifying the probability of resistance emergence and growth parameters

Once we determined the MIC for each strain in each antibiotic, we quantified the probability of emergence of resistance at the highest sub-MIC concentration, using a larger number of replicates for each strain and antibiotic combination (n=30–200 replicates per antibiotic; Table 1). For these measurements, we monitored growth in 96 well microplates as described above. We recorded growth or no growth outcomes based on both visible turbidity in cultures and whether the blank corrected OD_600_ value crossed 0.02 (the lower bound used by Curve Fitter software for growth curve analysis) (Delaney et al., 2013) or not. We estimated the probability of emergence of resistance as the ratio of replicates showing growth to the total number of replicates tested for each strain-antibiotic combination.

**Table 1.** Summary of sample sizes and outcomes for Experiment 2. The table shows the total number of replicate populations grown for each strain at its highest sub-MIC concentration in a given antibiotic, the number of replicates that showed visible growth, and the number of replicates used to extract growth parameters. Replicates in each successive column are a subset of the previous column.

| Antibiotic | Strain | Antibiotic concentration | Total replicates | Replicates with visible growth | Number of replicates used for growth parameter measurements |
| --- | --- | --- | --- | --- | --- |
| Strep | WT | 16 | 51 | 31 | 28 |
| Strep | Δnth-Δnei | 16 | 51 | 4 | 2 |
| Strep | ΔmutY | 16 | 45 | 21 | 6 |
| Strep | ΔmutS | 16 | 51 | 38 | 30 |
| Strep | ΔmutL | 16 | 51 | 38 | 30 |
| Strep | ΔmutH | 64 | 200 | 10 | 8 |
| Strep | ΔmutT | 1200 | 65 | 37 | 22 |
| Pol B | WT | 0.4 | 45 | 37 | 28 |
| Pol B | Δnth-Δnei | 1.6 | 30 | 5 | 5 |
| Pol B | ΔmutY | 3.2 | 30 | 9 | 9 |
| Pol B | ΔmutS | 3.2 | 30 | 6 | 6 |
| Pol B | ΔmutL | 0.8 | 45 | 17 | 17 |
| Pol B | ΔmutH | 0.4 | 45 | 42 | 29 |
| Pol B | ΔmutT | 0.4 | 45 | 43 | 29 |
| Cip | WT | 0.02 | 45 | 41 | 28 |
| Cip | Δnth-Δnei | 0.3 | 36 | 35 | 28 |
| Cip | ΔmutY | 0.04 | 45 | 23 | 11 |
| Cip | ΔmutS | 0.3 | 36 | 33 | 30 |
| Cip | ΔmutL | 0.08 | 45 | 28 | 14 |
| Cip | ΔmutH | 0.08 | 45 | 29 | 16 |
| Cip | ΔmutT | 0.08 | 50 | 26 | 7 |
| Cef | WT | 0.6 | 45 | 41 | 28 |
| Cef | Δnth-Δnei | 0.6 | 45 | 40 | 25 |
| Cef | ΔmutY | 0.8 | 45 | 18 | 5 |
| Cef | ΔmutS | 0.8 | 45 | 45 | 20 |
| Cef | ΔmutL | 0.8 | 45 | 40 | 29 |
| Cef | ΔmutH | 0.6 | 45 | 44 | 28 |
| Cef | ΔmutT | 0.8 | 45 | 31 | 26 |
| Rif | WT | 128 | 45 | 39 | 23 |
| Rif | Δnth-Δnei | 512 | 45 | 10 | 6 |
| Rif | ΔmutY | 2000 | 33 | 25 | 25 |
| Rif | ΔmutS | 2000 | 33 | 33 | 30 |
| Rif | ΔmutL | 2000 | 33 | 33 | 30 |
| Rif | ΔmutH | 2000 | 33 | 33 | 30 |
| Rif | ΔmutT | 2000 | 33 | 26 | 26 |
| Tet | WT | 1 | 45 | 7 | 3 |
| Tet | Δnth-Δnei | 1.6 | 30 | 24 | 24 |
| Tet | ΔmutY | 1 | 45 | 45 | 30 |
| Tet | ΔmutS | 1.6 | 30 | 30 | 30 |
| Tet | ΔmutL | 1.6 | 30 | 30 | 28 |
| Tet | ΔmutH | 1.6 | 30 | 30 | 30 |
| Tet | ΔmutT | 1 | 45 | 45 | 30 |

For a subset of the replicates showing growth above (Table 1), we tracked change in OD_600_ over time, and used these data to estimate final OD (indicator of biomass and magnitude of resistance), the lag time (indicating time to emergence of resistance mutations), and growth rate (indicating magnitude of resistance) for each strain-antibiotic-dose combination. We estimated growth rate using the Curve Fitter software (Delaney et al., 2013), which fits a linear regression to log(OD) vs. time data. We estimated lag time as the time to reach OD=0.02, which represents the lowest threshold of exponential growth detection of the Curve Fitter software; and final OD as the observed OD after 18 h of growth. We also estimated growth rate and final OD of each strain and antibiotic from the lower antibiotic concentrations used in Experiment 1 (Fig. S4 and S5)

### Sanger sequencing to identify mutations in antibiotic resistance genes

We performed Sanger sequencing of target genes for each antibiotic (Table S1) to identify single nucleotide substitutions observed after 18 h of growth, from the high-replication growth assays at sub-MIC concentrations described above (Experiment 2). We randomly chose 6 replicate wells showing growth, made cryostocks, and streaked each culture on LB agar plates incubated at 37 °C overnight. From each plate, we randomly chose a single colony (i.e., one clone per replicate population), performed colony PCR to amplify the respective target gene(s) (Table S1), and checked the PCR products using gel electrophoresis. We purified amplicons using magnetic beads in 96 well plates, thoroughly mixing 20 µL PCR product with 20 µL beads and incubating at room temperature for 5 min. We placed the plate on a magnetic stand for 5 min to allow DNA to be adsorbed onto the beads. We removed the supernatant, washed the beads with 80% ethanol twice, and eluted DNA by adding 20 µL Tris-HCl buffer. We Sanger-sequenced PCR products using the same primers used for amplification (Table S1). Sanger sequencing of target genes was also performed using the same method for a few strains at lower antibiotic concentrations (Table S2)

### Whole genome sequencing

To identify non-target gene mutations, we randomly chose 3 replicate populations showing growth in Experiment 2, for population level Whole Genome Sequencing. We also sequenced the ancestral population of each strain to identify background mutations in each strain (apart from the intended DNA repair gene deletion). We inoculated 30 µL of each cryostock in 1 mL LB in 15 mL falcon centrifuge tubes (Tarson), and incubated at 37 °C with orbital shaking at 200 rpm until we observed turbid growth (typically 4 h, except Strep and Tet, which required 7-8 h). We extracted genomic DNA using the Qiagen DNeasy Blood & Tissue Kit, following kit instructions. We quantified DNA concentration using Qubit for a subset of 2 randomly chosen samples per antibiotic. We prepared paired-end libraries for each population using the Illumina DNA Prep (M) Tagmentation kit (CAT No: 20060059) or Ilumina Nextera XT DNA Library Preparation Kit for 96 samples (CAT No: FC-131-1096). Sequencing was performed on the NovaSeq 6000 platform (2×100 paired-end or 2X150 paired-end), with an average depth of ∼156X per sample (depth range ∼80–242X; Table S3). We first trimmed adapters (CTGTCTCTTATACACATCT) using fastp version 0.23.4 (Chen et al., 2018), and aligned reads to the *E. coli* reference genome (Refseq: GCF_000005845.2) to identify mutations using the breseq v.0.39.0 pipeline (Deatherage and Barrick, 2014). We filtered out mutations present in the respective ancestor of each strain, to identify new mutations that arose under antibiotic stress. We confirmed that three mutations present in our WT ancestor (Sane et al., 2025) were found at 100% frequency in all samples, as expected.

### Experiment 3: Measuring the cost of resistance mutations

Next, we measured the cost of resistance mutations that emerge in the first 18 hours of antibiotic exposure. We conducted a new experiment, inoculating 6 replicate wells at the highest sub-MIC concentration (as described for Experiment 2) for each strain-antibiotic combination, and then propagating cultures with putative resistance mutations for one more growth cycle of 18 h in LB media without antibiotic (1:100 dilution). We measured the final OD and growth rate for both cycles. We measured the cost of resistance mutations as: final OD or growth rate in the second growth cycle - final OD or growth rate of the respective ancestor grown in LB without prior exposure to antibiotic (control data from Experiment 1). Positive values would indicate that the resistance mutations that emerged after antibiotic exposure of 18 hours were beneficial even in the absence of the antibiotic (i.e., they increased growth in LB compared to the ancestor). Conversely, negative values would indicate that the mutations were costly in the absence of antibiotic, with the absolute parameter value indicating the magnitude of the cost. No difference would indicate neutral mutations.

Sometimes — e.g., *Δnth-Δnei* and *ΔmutS* in Pol B — none of the 6 replicate wells in this experiment showed any growth during the first cycle; this was not unexpected given the low probability of resistance. In such cases, we repeated the experiment until we obtained data for at least one replicate with growth in cycle 1. For some of these cases, we could not conduct statistical analysis given low replication; these data points are marked accordingly in Fig. 3 and S6.

### Data analysis

We used R version 4.3.3 for all data analysis (R Development Core Team, 2024). To compare the probability of observing growth (indicating resistance) of different mutators (Experiment 2), we performed pairwise Chi-square tests. To compare growth parameters of mutators grown in the same sub-MIC concentration (lag time, final OD and growth rate), we performed pairwise Welch’s t-tests (Experiment 2). For comparisons among multiple mutators growing in the same concentration of an antibiotic, we used Benjamini-Hochberg tests to correct for multiple comparisons. To evaluate the cost of resistance mutations (Experiment 3), we tested for differences in growth rate or final OD using t tests.

## RESULTS

### Specific mutation bias — not mutation rate — predicts survival and growth under several antibiotics

We first determined the MIC (minimum inhibitory concentration) for each mutator strain and WT in each of six antibiotics, using a small number of replicate cultures (n=3; Fig. 1A) (Experiment 1). As expected, in most cases the WT had a lower MIC than mutators, with two exceptions. In Strep, WT had a higher MIC than *ΔmutY*, and in Pol B *ΔmutH* and *ΔmutT* had similar MIC as WT (Fig. 2A; see Fig. S2 for raw growth curves). In Strep and Pol B, *ΔmutT* and *ΔmutY* respectively had much higher MICs than other mutators; but in other antibiotics, no single strain clearly outperformed the others (Fig. 2A). Notably, the MIC values were not consistently higher for strains with the highest mutation rates. We checked for the presence of antibiotics in the media by inoculating fresh culture of the strains in spent media from experiment 2 after 18 hours of antibiotic exposure. Growth was still suppressed compared to the control cultures where fresh cultures of strains were inoculated in spent LB media without any antibiotics in most cases. This suggests that antibiotic was still present in the media at the endpoint of our experiments. (Fig. S3).

**Fig. 2.**
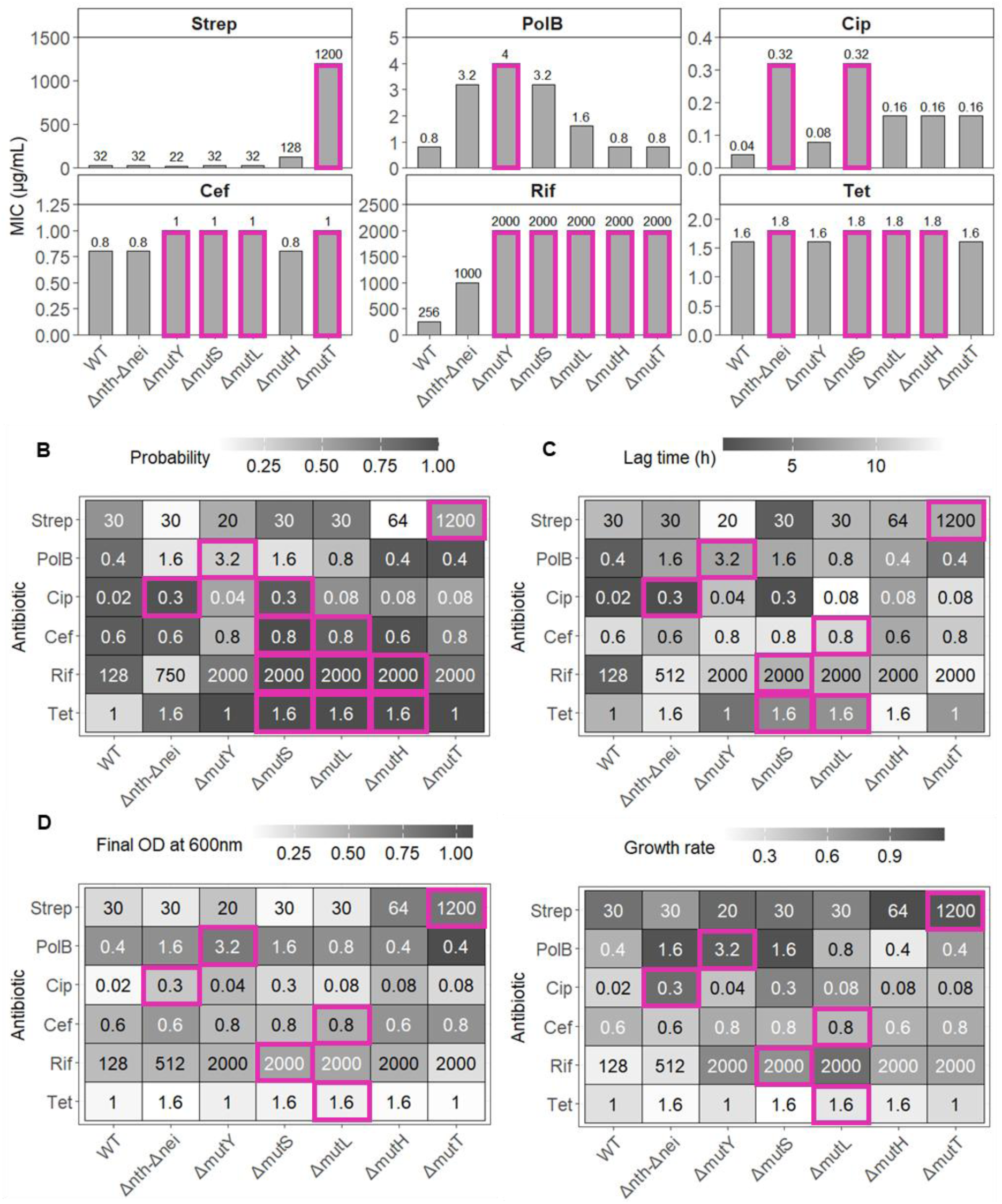
Survival and growth parameters of strains in each antibiotic. **(A)** Bar plots showing MIC values (µg/ml) inferred from Experiment 1, also indicated on top of bars for clarity. Pink boxes indicate strains with the highest MIC in each antibiotic. **(B)** Heatmap showing the probability of resistance emergence for each strain-antibiotic concentration estimated from Experiment 2. Sample sizes are given in Table 2. **(C–E)** Heatmaps showing growth parameters estimated from growth curves obtained in Experiment 2. Sample sizes (number of replicates that showed growth) are given in Table 2. **(C)** Lag time (h) **(D)** Final OD_600_ **(E)** Growth rate (h^-1^). In panels B–E, numbers inside cells indicate the highest sub-MIC concentration at which the resistance probability or growth parameter was estimated. Pink boxes indicate the fittest mutators according to the specific parameter, in each antibiotic.

**Table 2.**
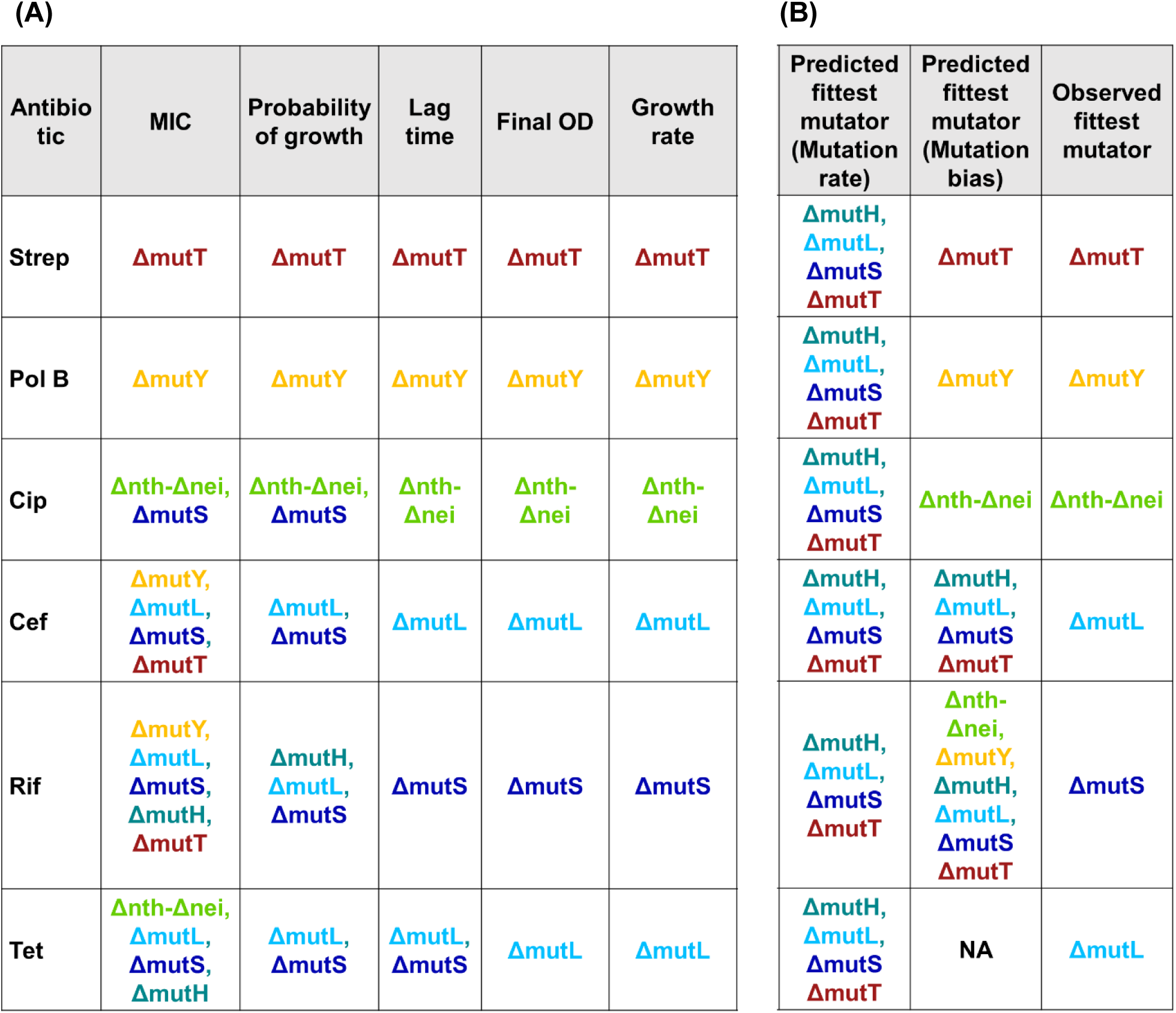
(A) Fittest mutators identified after measurement of each fitness parameter, (B) Comparison of the observed fittest mutator with the predicted fittest mutator according to mutation rate and mutation bias.

Though informative, MIC values do not provide a quantitative metric to predict the likelihood and time to emergence of resistance. Hence, we measured the probability of resistance emerging at the highest sub-MIC concentration of each antibiotic (identified from the MIC assays above), by estimating the proportion of independent populations that showed detectable growth across a large set of replicates of each strain (n>=30; Fig. 1B; see Fig. S7 for growth curves) (Experiment 2). The results broadly corroborated the MIC outcomes, and also allowed us to distinguish between strains that had identical MIC values. For instance, of the four strains with similar MIC values for Cef, *ΔmutL* and *ΔmutS* had higher probability of growth than *ΔmutT* and *ΔmutY* at a concentration of 0.8 µg/mL antibiotic (Fig. 2B, Table S4A). However, in Rif and Tet, many strains had similar survival at a very high antibiotic concentration, and we could still not distinguish between these (Table S4A). As a third proxy for antibiotic survival, we next estimated lag time from the growth measurements: a shorter lag time indicates faster emergence of resistance mutations. Again, the results broadly mirrored survival as measured by MIC and probability of growth for Strep and Pol B, but revealed an advantage for *Δnth-Δnei* relative to *ΔmutS* in Cip, and for *ΔmutL* over *ΔmutS* in Cef (Fig. 2C; Fig. S7, Table S4B). In Rif and Tet, lag time also distinguished *ΔmutL* (Tet) and *ΔmutS* (Rif and Tet) as the fittest mutators (Table S4B). Finally, we estimated the magnitude of resistance using growth rate and final OD from the above data. Although neither mutation bias nor rate directly predict these parameters, these two parameters resolved *ΔmutL* as the fittest mutator in Tet (Fig. 2D and 2E, Table S4B). Again, different strains with the same mutation rate showed variable performance in each antibiotic.

A summary of all the survival and growth metrics (Table 2) highlights that mutation rate is a poor predictor of antibiotic survival overall. On the other hand, mutation bias (i.e., higher mutation rates for specific types of mutations) accurately predicted which strain was more likely to rapidly and/or consistently give rise to resistance in three antibiotics: Strep, Pol B, and Cip (Table 2). In particular, in both Pol B and Cip, mutators with intermediate mutation rate showed greater resistance than mutators with the highest mutation rate. However, in Cef, Rif and Tet, neither mutation rate nor mutation bias clearly explained the high fitness of *ΔmutL* or *ΔmutS*, which outperformed other strains with a similar mutation rate (*ΔmutH* and *ΔmutT*) or a similar mutation spectrum (*ΔmutH*). Cef is an interesting case because there are many reported beneficial resistance mutations for different target genes; hence, all strains could rapidly sample different beneficial mutations. For resistance via *ftsI, ΔmutT* was predicted to be the fittest mutator; for resistance via *marR* (efflux), several strains have appropriate mutation spectra, including *ΔmutL* (Fig. 1D). Hence, the reason for the “best” performance of *ΔmutL* is unclear; but since *ΔmutT* was not the fittest strain in Cef, we predicted that Cef resistance in our experiment likely involved *marR* mutations. Note that Rif and Tet are special cases for which we did not have a clear prediction about which specific strain would be more likely to sample resistance mutations. For Rif, there are diverse paths to resistance, and each strain had the potential to rapidly sample at least one type of resistance mutation (Fig. 1D). On the other hand, our *E. coli* strains lack clear target genes for Tet resistance. Hence, mutation bias is not informative or predictive for either Rif or Tet; and the reasons for the high survival of *ΔmutS* and *ΔmutL* in these antibiotics is unclear. Overall, of the 4 antibiotics with a clear prediction for highest survival and growth based on mutation bias, results for 3 antibiotics were consistent with our predictions. In contrast, in no case did mutation rate accurately predict the identity of mutator(s) with highest fitness. Thus, increasing mutation rate relative to the WT is broadly beneficial, as shown in prior studies; but beyond a 10x increase in mutation rate (intermediate mutation rate), there is no further advantage.

### Resistance is generally costly in the absence of antibiotics

We next examined the fitness cost of resistance mutations that arose during the first 18 h of antibiotic exposure. By comparing the growth rate and final OD of putatively resistant populations propagated in antibiotic-free medium to those of their respective ancestors (also grown in antibiotic-free LB), we estimated the cost of resistance. In most cases, resistance was associated with a reduction in fitness, demonstrating that resistance evolution is generally costly in the absence of antibiotics (Fig. 3 and S6). Prominent exceptions were Rif and Pol B, in which 3-4 strains evolved resistance without significant costs. In addition, in 5 of 6 antibiotics, resistance evolved in *ΔmutL* was not costly (Fig. 3). Notably, very few of the resistant isolates exhibited higher fitness than their corresponding ancestors, indicating an absence of broadly beneficial resistance mutations under our assay conditions. Finally, Tet resistance was generally most costly in terms of growth rate (Fig. 3). Together, these results suggest that apart from *ΔmutL,* all mutators evolve costly resistance across antibiotics.

**Fig. 3:**
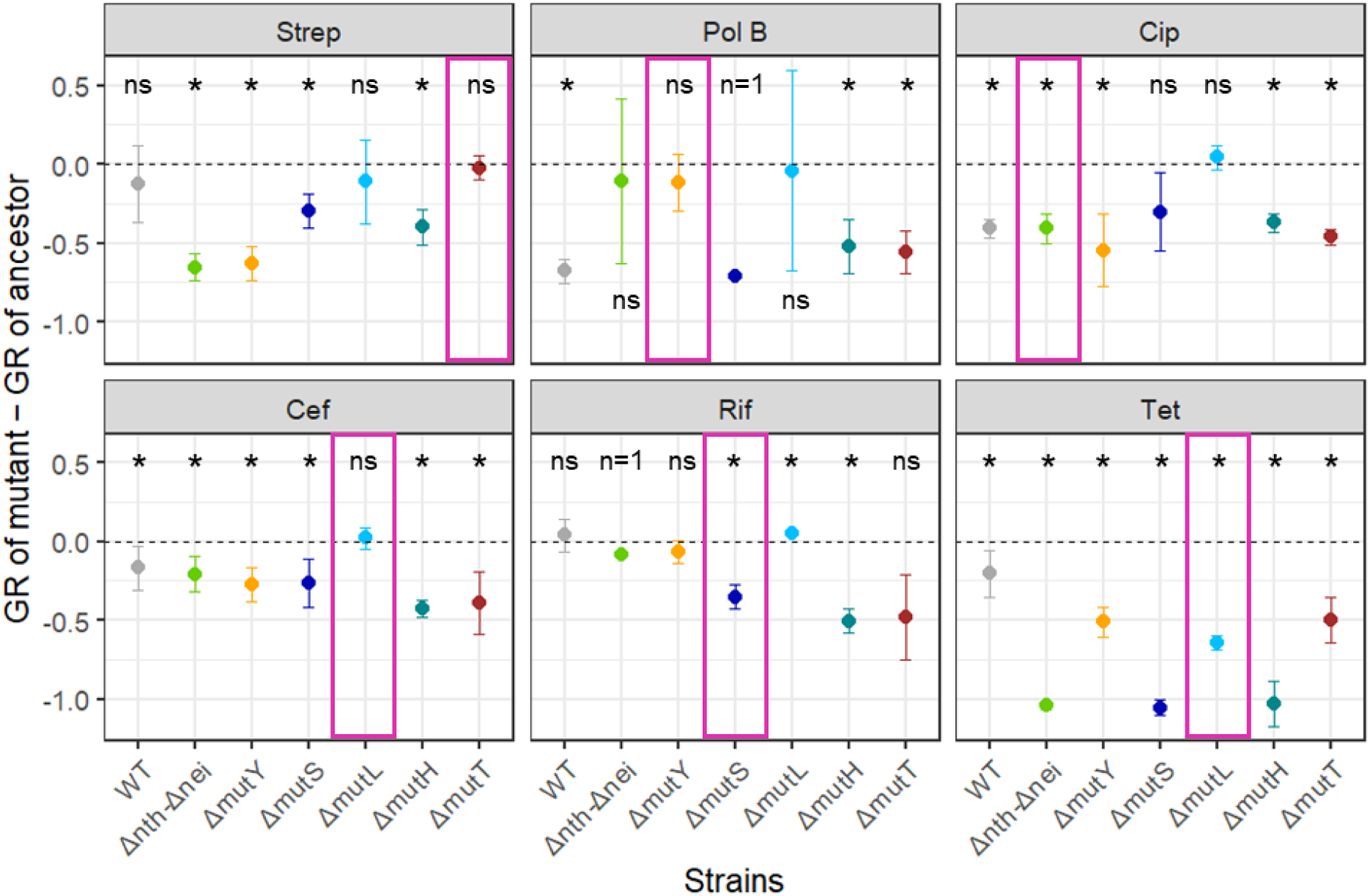
Cost of antibiotic resistance mutations in absence of antibiotic. Cost was calculated as the difference between the growth rate (GR) of resistant populations and that of their corresponding ancestral strains, both grown in antibiotic-free LB. Points represent differences in growth rate between resistant mutants and respective ancestors (n=2-6, mean ± standard deviation). In two cases, only one replicate showed measurable growth (i.e., showed resistance); these are marked, and were not analysed statistically. The dashed horizontal line denotes no difference from the ancestor (i.e., no cost). Negative values indicate that resistance mutations reduced growth rate relative to the ancestor (fitness cost), whereas positive values indicate a benefit in the absence of antibiotic. Asterisks indicate strain-antibiotic combinations for which costs were significantly different from zero (t-tests, p < 0.05).

### Resistance mutations reflect strain-specific mutation spectra, with notable exceptions

If specific mutation biases explain the emergence of resistance, resistance mutations should reflect strain-specific mutation spectra inferred from mutation accumulation experiments (i.e., under genetic drift; Sane et al., 2025). To test this, we randomly chose three replicates from showing growth at the highest sub-MIC concentration for each antibiotic, and sequenced the whole population using NGS. As predicted, mutations in known resistance genes closely matched the mutation spectrum of each strain (Fig. 4), further supported by Sanger sequencing of target genes from more replicates (Fig. S8). We observed similar results from NGS and Sanger sequencing at intermediate antibiotic concentrations (Table S2 and S5). Importantly, we found mutations in target genes of some antibiotics at low and intermediate antibiotic concentrations using Sanger sequencing of a few strains (Table S3). These mutations also matched the mutation spectra for the respective strains. However, in the lowest concentrations tested using Sanger sequencing, such mutations were rare (only one replicate). Note that not all mutation types that dominate a strain’s mutation spectrum are observed in our experiments, because only those mutations that confer resistance can rise to sufficiently high frequencies. For instance, the mutation spectra of *ΔmutH*, *ΔmutL* and *ΔmutS* consist of both CG◊TA and TA◊CG mutations (Fig. 1D); but in Cip, only the former mutations in *gyrA* confer high resistance frequently and were observed in our dataset (Fig. 4). Thus, we observed a strong signature of each strain’s mutation spectrum in the set of putative mutations responsible for resistance to high doses of antibiotics.

**Fig. 4:**
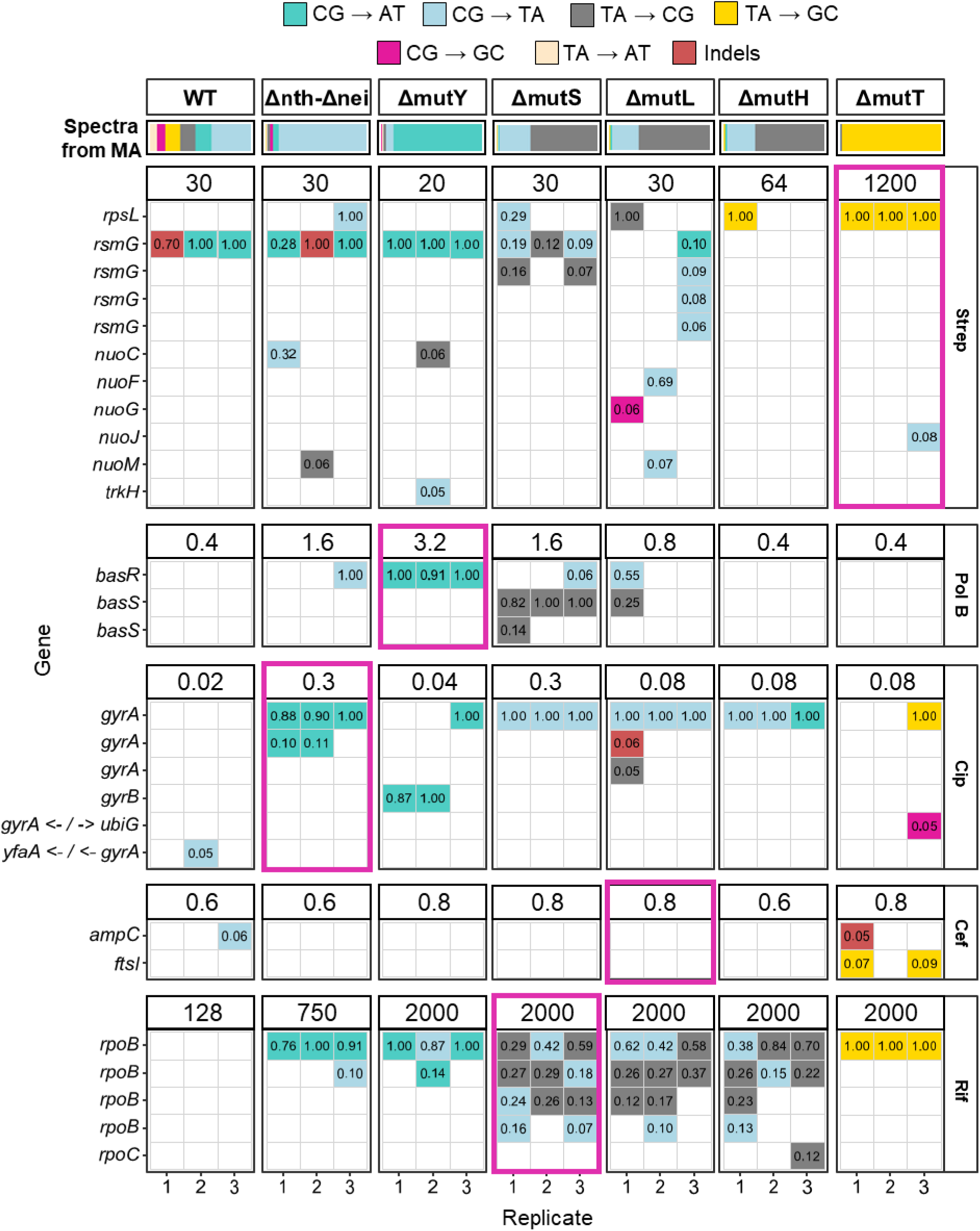
The spectrum of target gene mutations observed in resistant populations of mutators in each antibiotic. At the top, the mutation spectrum of each strain is indicated; this is identical to Fig. 1D. For each antibiotic (indicated on the far right), the occurrence of mutations in known target genes is shown, for each of three replicate populations per strain with NGS data (represented in columns). Numbers above each panel indicate the highest sub-MIC concentration (µg/mL) at which the strain was tested (Experiment 2). Each row indicates a distinct mutation in the gene indicated on the left. Repeated gene names for a given strain and antibiotic combination indicate mutations observed at different positions in the same gene. However, across strains, the row identity is not meaningful. E.g., in Strep, several strains have *rpsL* mutations (first row); these are not necessarily at the same position across strains. Numbers inside each cell indicate the frequency at which the mutation was observed (a value of 1 indicates that the allele was fixed in the population). Pink boxes around panels indicate the fittest mutator in each antibiotic.

An interesting exception was a *ΔmutH* population in Strep, which showed an TA→GC mutation that is atypical of its mutation spectrum (Fig. 4). However, this particular population also had a (C_6◊7_) mutation in the *mutT* gene at 100% frequency, known to be a loss-of-function mutation (Elgrail et al., 2024). Thus, the *mutT* loss-of-function mutation likely arose first, which facilitated the sampling of the TA→GC resistance mutation that is also observed repeatedly in *ΔmutT* populations. We also observed several CG→AT mutations in *Δnth-Δnei* in Strep, Cip, and Rif, whereas only 5.6% of mutations are expected to be CG→AT. Most (86.4%) of mutations in this strain should be CG→TA, which includes known resistance alleles. Unlike the *ΔmutH* population discussed above, neither the ancestral *Δnth-Δnei* strain nor the resistant populations had any mutations associated with known DNA repair genes that could account for the unexpected mutation bias. In Rif and Strep as well, *Δnth-Δnei* sampled the rare type of mutation, but in those cases the strain performed poorly. However, at 0.3 µg/mL Cip, *Δnth-Δnei* was clearly the fittest strain, acquiring two *gyrA* mutations each (both CG→AT), in two of three replicates (Fig. 4). Notably, at a relatively lower Cip concentration (0.16 µg/mL) — still very high, as reflected by the very low probability of growth in other strains (Fig. 2B) — we did observe the expected CG→TA resistance mutations in *gyrA* (Table S5). Thus, as selection strength increases, relatively rare mutations can evidently outcompete more commonly sampled mutational types.

In most antibiotics, all replicates of strains with high survival at high concentrations showed a target resistance mutation (Fig. 4 and 5). In several cases we did not observe any known target gene mutations, though most of these also showed very poor performance (e.g., *ΔmutT* and *ΔmutH* in Pol B, Fig. 5). Using the NGS data, we therefore categorized all non-target mutations that could potentially confer antibiotic resistance (Fig. 5). As predicted by the phenotypic data, nearly all resistant populations in Cef had efflux pump mutations, including in *marR* (Fig. S9). Hence, we asked: do efflux pump gene mutations also reflect strain-specific mutational biases? The most prevalent known *marR* mutations for Cef resistance are TA→AT mutations (Comprehensive Antibiotic Resistance Database (CARD) (Alcock et al., 2023); MicroBIGG-E (Feldgarden et al., 2021)); but we did not observe such mutations in any strain, likely because this mutation type is rare in all our strains (Fig. 1D). CG→TA mutations are also frequently reported to drive resistance to Cef; these were observed in *ΔmutH, ΔmutL* and *ΔmutS*, matching their respective mutation spectra (Fig. 1D and S9). Presumably, the specific mutations sampled by *ΔmutL* conferred higher resistance than those sampled by other strains, driving its high performance in Cef. Finally, in Strep, two replicates of *ΔmutH* stand out: they showed good growth at high concentrations (64 µg/mL), but did not have any high frequency mutations in any genes (Fig. 5), and it remains unclear how these specific populations achieved this resistance. Note that only 5 out of 200 tested replicates of *ΔmutH* showed any growth at this Strep concentration, implying that resistance in general is not frequent.

**Fig. 5:**
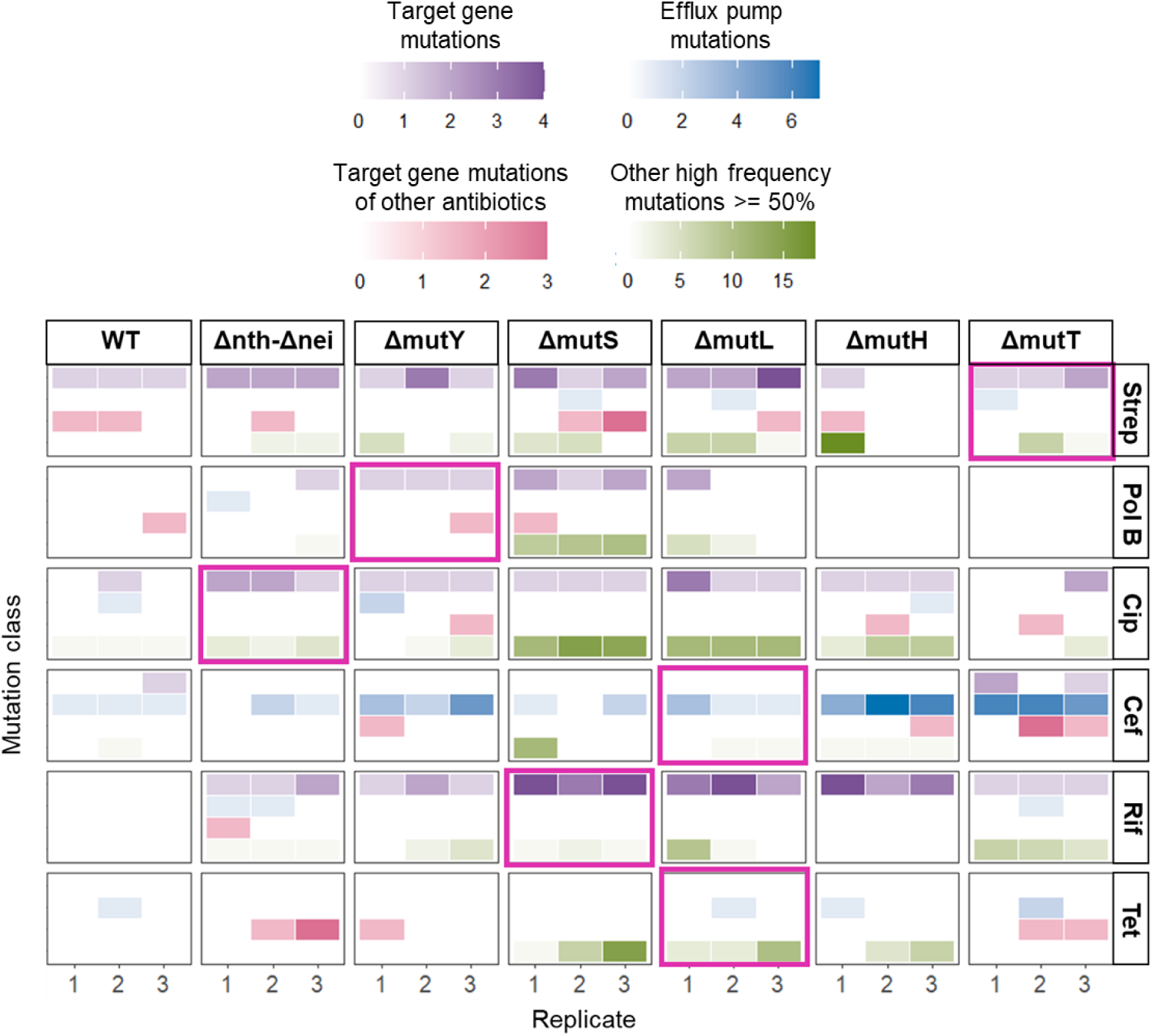
Number of mutations reported from whole genome sequencing in different mutation categories. Heatmaps show the number of mutations detected by whole-genome sequencing in resistant populations evolved under six antibiotics (rows) across wild-type (WT) and mutator backgrounds (columns). Within each strain-antibiotic combination, mutations are grouped into four categories: mutations in known antibiotic target genes, mutations in efflux pump-associated genes, mutations in resistance genes associated with other antibiotics, and any other high-frequency mutations (>50% frequency within a population). Columns labelled 1–3 in the X axis represent replicate resistant populations. Colour intensity of the tiles indicates the number of mutations detected in each category (colour key at the top). Pink boxes highlight the strain that exhibited the highest fitness in each antibiotic.

Three important broader points emerge from this analysis of genome-wide mutations. First, uniquely for Cef, resistance was generally associated with efflux pump mutations across strains and antibiotic concentrations. Second, *ΔmutL* and *ΔmutS* acquired several high frequency mutations in non-target genes across antibiotics (Fig. 5), potentially explaining their high performance in Cef, Rif and Tet. Similarly, mutations in efflux pump genes may explain high resistance of *ΔmutL* in Tet. More generally, across all antibiotics and strains, we often observed mutations in target genes of other antibiotics to which the populations were not exposed in our experiment, at frequencies ranging from 5-15% (Fig. S10). Mutations in *rpoB* and *rpoC* were most common within this dataset. Third, as expected, the spectrum of high frequency mutations (at least 80%) typically matched the spectrum obtained from mutation accumulation experiments for each strain (Fig. S11A and S12). Additionally, *ΔmutS* and *ΔmutL* in all antibiotics, and *ΔmutH* and WT in Cip sampled many high frequency indels, again consistent with their mutation spectra under genetic drift (Fig. S11B). However, unexpectedly, in Strep, Cip and Rif, *Δnth-Δnei* sampled many high frequency CG→AT mutations, which should be rare in this strain (Fig. S12).

## DISCUSSION

Antibiotic resistance mutations play a central role in the emergence and persistence of resistant bacterial populations. These mutations — which often alter antibiotic targets, reduce drug uptake, or enhance efflux — provide the initial genetic variation upon which selection acts (Martínez & Baquero, 2000; Blair et al., 2015). Here, we show that the supply rate of beneficial mutations under strong antibiotic selection depends on the rate of specific types of mutations which happen to confer high level resistance (i.e., the mutation bias). In this study, we did not see a strong effect of mutation rate on mutator survival under antibiotics such as Cip and Pol B, where mutators with intermediate mutation rate (*Δnth-Δnei* and *ΔmutY*) outperformed strains with the highest mutation rate. However, the rank order of survival was well predicted by the strain-specific mutation biases (Table 2). Additionally, mutators with similarly high mutation rates (i.e., *ΔmutT, ΔmutH, ΔmutL,* and *ΔmutS*) did not show similar fitness in any antibiotic, as would be expected if mutation rate alone determined the emergence of resistance. Notably, the same high-rate mutators were not the fittest across all tested antibiotics. Thus, mutation rate alone has poor predictive power for rapid emergence of resistance. Instead, along with prior studies (Couce et al., 2013; Couce et al., 2015; Barber and Couce., 2025; Payne et al., 2019), our results demonstrate the importance of specific mutation bias in driving rapid antibiotic resistance. For instance, in fluctuation tests on agar plates, *ΔmutT* had more colonies with higher growth rate than *ΔmutY* in Streptomycin; whereas in Rifampicin (which belongs to the same class of antibiotics as Rifamycin, used in our study) the opposite was observed (Couce et al., 2013). Both results are consistent with our data, though in Rif we identified other strains with even faster emergence and higher magnitude of resistance than either *ΔmutY* and or *ΔmutT*. In Tetracycline, Couce and colleagues found that both *ΔmutY* and *ΔmutT* performed similarly, as was also observed in our results. In another study, after 28 days of evolution in increasing concentrations of Cefotaxime, *ΔmutH* populations showed a significant drop in survival at concentrations where *ΔmutT* could still survive, associated with more resistance mutations in both genomic and plasmid borne target genes (Couce et al., 2015). Importantly, as we observed here for *ΔmutT*, resistance mutations matched the strain-specific mutation spectra, although in their study resistance occurred via mutations in the target gene *ftsI* (more TA→GC mutations in *ΔmutT* populations, and more CG◊TA and TA◊CG mutations in *ΔmutH*). Together, these studies show that mutation bias is a stronger predictor of emergence of antibiotic resistance mutations than mutation rate.

In contrast to these studies whose results corroborate ours, there are significant departures in some cases. For instance, Shibai and colleagues evolved multiple mutators including *ΔmutT, ΔmutH, ΔmutL* and *ΔmutS* in Ciprofloxacin. They reported similar performance for all these strains, contrary to our results where *mutS* had higher MIC in Ciprofloxacin (Shibai et al., 2025). However, theirs was a longer-term evolution study, allowing mutators with high mutation rates to sample many different beneficial mutations, which also lead to a doubling of the ancestral MIC values. Whether their strains sampled different types of first-step resistance mutations, and whether they aligned with the respective mutation spectra, remains unknown. Another study by Miller and colleagues measured the MIC of *ΔmutY, ΔmutT, ΔmutH, ΔmutL* and *ΔmutS* in rifampicin, ciprofloxacin, cefotaxime and polymyxin B (Miller et al., 2002). Similar to our results, all these strains had the same MIC in rifampicin. However, both *ΔmutS* and *ΔmutH* had similarly high MICs in ciprofloxacin; *ΔmutT, ΔmutL* and *ΔmutS* performed best in cefotaxime; and *ΔmutS* and the ancestral strain had higher MICs than the other mutator strains in polymyxin B. These results differ from ours (Fig. 2A), potentially due to several methodological differences that can alter MIC values, e.g., nutrient media (LB vs. Mueller Hinton Agar), the use of liquid vs. solid media, and distinct strain backgrounds. We hope that future work with various strains and bacterial species with diverse spectra can broaden our understanding of the effects of mutation bias on antibiotic resistance.

Next, we discuss some unexpected results that present interesting avenues for further work. First, in some cases the mutation spectra of resistant populations did not match the spectra expected from mutation accumulation studies (Fig. S13). For instance, *ΔmutY* and *ΔmutT* are strongly transversion biased strains, yet they sampled many low frequency (<30%) transition mutations. Notably, the high frequency mutations in these cases were usually transversions, as expected. Hence, we speculate that the low-frequency transitions are gradually repaired in these populations, and do not reach high frequencies. We also observed unexpected mutation biases in Cip, Strep and Rif, where *Δnth-Δnei* populations often sampled CG→AT mutations that are rare in its mutation spectrum. In Cip, as discussed in the Results section, we speculate that a gene x environment interaction leads to extremely rare CG→ AT mutations being beneficial (and therefore being selected) at high Cip concentrations, whereas the more frequently sampled CG→TA mutations are sufficient to survive a lower concentration. Thus, this represents an interesting case where strong selection overcomes constraints placed by arrival (mutation) bias. Similar effects may also explain the high-frequency rare mutation types (CG→AT) in Strep and Rif; but this requires verification. Second, *ΔmutL* and *ΔmutS* emerged as the fittest mutators in Rif, Cef, and Tet, and the reasons are not clear. Notably, *ΔmutH*, which is part of the same repair pathway, does not show a similar advantage. We speculate that the unique resistance outcomes for *ΔmutS* and *ΔmutL* are related to the mechanism of repair that these two proteins perform, which is distinct from MutH. Given that loss of function of *mutL* and *mutS* is observed often in clinical antibiotic resistance (Hall et al., 2006; Boyce, 2022), we suggest that our results reflect a broader phenomenon, with fundamental underlying mechanisms that should be explored further. Finally, the two replicates of *ΔmutH* that grew and survived in 64 µg/ml Strep, without acquiring any detectable mutations, suggests a role for physiological mechanisms of resistance. All these unexpected observations are fertile grounds for future work.

Our results also have important implications for understanding the genetic basis of adaptation. Prior theoretical and empirical work shows that reversing the ancestral mutation bias should facilitate adaptation in new environments (Sane et al., 2023; Tuffaha et al., 2023; Sane et al., 2025; Parveen et al., 2025). Our current results demonstrate conditions when this advantage does not hold. Wild type *E. coli* has a transition-biased mutation spectrum, so we had predicted that transversion-biased mutators (here, *ΔmutT and ΔmutY*) should have a broad adaptive advantage. Clearly, this is not the case under antibiotic selection. The broad advantage of a bias reversal stems from increased sampling of previously poorly explored types of mutations, in conditions where beneficial mutations are limited. However, under strong antibiotic selection, the set of beneficial mutations consists of specific types of mutations in specific target genes; depending on the antibiotic, the total number of possible beneficial mutations varies. But because resistance mutations are often costly, they are not already fixed in populations, and thus can arise anew and fix under high antibiotic concentrations. Hence, adaptation to antibiotics is limited by a population’s ability to sample these specific types of mutations rather than a general depletion of beneficial mutations. Thus, our results demonstrate boundary conditions for the advantage of bias reversals during adaptation. We expect that bias reversals will similarly fail to predict evolutionary outcomes in other selection regimes where adaptation is not limited by a broad depletion of beneficial mutations.

In conclusion, our work contributes to the growing understanding of the importance of specific mutation biases in driving drug resistance and disease outcomes, including in various cancers (Leighow et al., 2020; Tuffaha et al., 2025), and opens several avenues for further work. Note that although we focused here on genetic changes leading to altered mutation spectra (via deletion of DNA repair genes), mutation biases can also be induced by environmental factors such as the niche of the bacteria (Ruis et al., 2023), high temperature (Shewaramani et al., 2017) and nutrient limitation and availability (Maharjan and Ferenci, 2017, Gifford et al., 2024). Therefore, understanding and quantifying the incidence and magnitude of mutation biases should be broadly useful to predict antibiotic resistance under diverse contexts. However, it remains to be seen whether the influence of mutation bias is relevant in the long-term. We speculate that the observed short-term advantage of specific mutators in specific antibiotics will weaken over time, due to 2^nd^ or 3^rd^ step mutations that open alternative paths of resistance, or compensatory mutations that mitigate the cost of resistance. Additionally, mutators rapidly accumulate deleterious mutations, which may alter their survival and competitive ability. Testing whether short-term advantages observed in the lab persist long-term is thus important, especially in the clinic. Further work could also determine the mechanism through which some of our (previously unknown) putative resistance mutations may act; e.g., CG→AT mutations sampled by *Δnth-Δnei* in Cip. We hope that such studies will improve our ability to predict the emergence and effects of drug resistant mutations.

## DATA AVAILABILITY

All data underlying this article are available in the article and in its online supplementary material.

## ACKNOWLEDGEMENTS

We thank Parth Raval and members of the Agashe lab for discussion and critical comments on the manuscript. We acknowledge funding and support from the National Centre for Biological Sciences; the DBT/Wellcome Trust India Alliance (grant IA/S/23/2/506989 to DA); and a University Grants Commission Fellowship to AC.

## AUTHOR CONTRIBUTIONS

AC and DA conceived and designed the project; AC conducted experiments and analysed data; DA directed all work and analyses; AC and DA wrote the manuscript; DA acquired funding.

## Notes

### Competing Interest Statement

The authors have declared no competing interest.

## References

Alcock, B. P., Huynh, W., Chalil, R., Smith, K. W., Raphenya, A. R., Wlodarski, M. A., Edalatmand, A., Petkau, A., Syed, S. A., Tsang, K. K., Baker, S. J. C., Dave, M., McCarthy, M. C., Mukiri, K. M., Nasir, J. A., Golbon, B., Imtiaz, H., Jiang, X., Kaur, K.,…McArthur, A. G. (2023). CARD 2023: expanded curation, support for machine learning, and resistome prediction at the Comprehensive Antibiotic Resistance Database. Nucleic Acids Research, 51(D1), D690–D699. 10.1093/nar/gkac920

Barber, J. N., & Couce, A. (2025). Mutation-biased adaptation is consequential even in large bacterial populations. 10.1101/2025.06.16.655099

Blair, J. M. A., Webber, M. A., Baylay, A. J., Ogbolu, D. O., & Piddock, L. J. v. (2015). Molecular mechanisms of antibiotic resistance. Nature Reviews Microbiology, 13(1), 42–51. 10.1038/nrmicro3380

Boyce, K. J. (2022). Mutators Enhance Adaptive Micro-Evolution in Pathogenic Microbes. Microorganisms, 10(2), 442. 10.3390/microorganisms10020442

Cano, A. v., Rozhoňová, H., Stoltzfus, A., McCandlish, D. M., & Payne, J. L. (2022). Mutation bias shapes the spectrum of adaptive substitutions. Proceedings of the National Academy of Sciences, 119(7). 10.1073/pnas.2119720119

Chen, S., Zhou, Y., Chen, Y., & Gu, J. (2018). fastp: an ultra-fast all-in-one FASTQ preprocessor. Bioinformatics, 34(17), i884–i890. 10.1093/bioinformatics/bty560

Chowdhury, N., Suhani, S., Purkaystha, A., Begum, M. K., Raihan, T., Alam, Md. J., Islam, K., & Azad, A. K. (2019). Identification of *AcrAB-TolC* Efflux Pump Genes and Detection of Mutation in Efflux Repressor *AcrR* from Omeprazole Responsive Multidrug-Resistant *Escherichia coli* Isolates Causing Urinary Tract Infections. Microbiology Insights, 12. 10.1177/1178636119889629

Couce, A., Guelfo, J. R., & Blázquez, J. (2013). Mutational Spectrum Drives the Rise of Mutator Bacteria. PLoS Genetics, 9(1), e1003167. 10.1371/journal.pgen.1003167

Couce, A., Rodríguez-Rojas, A., & Blázquez, J. (2015). Bypass of genetic constraints during mutator evolution to antibiotic resistance. Proceedings of the Royal Society B: Biological Sciences, 282(1804), 20142698. 10.1098/rspb.2014.2698

Deatherage, D. E., & Barrick, J. E. (2014). Identification of Mutations in Laboratory-Evolved Microbes from Next-Generation Sequencing Data Using breseq (pp. 165–188). 10.1007/978-1-4939-0554-6_12

Delaney, N. F., Kaczmarek, M. E., Ward, L. M., Swanson, P. K., Lee, M.-C., & Marx, C. J. (2013). Development of an optimized medium, strain and high-throughput culturing methods for Methylobacterium extorquens. PloS One, 8(4), e62957. 10.1371/journal.pone.0062957

Elgrail, M. M., Sprouffske, K., Dartey, J. O., & Garcia, A. M. (2024). Emergence of a multilocus mutator genotype in mutator *Escherichia coli* experimental populations under repeated lethal selection. Journal of Evolutionary Biology, 37(3), 346–352. 10.1093/jeb/voae007

Feldgarden, M., Brover, V., Gonzalez-Escalona, N., Frye, J. G., Haendiges, J., Haft, D. H., Hoffmann, M., Pettengill, J. B., Prasad, A. B., Tillman, G. E., Tyson, G. H., & Klimke, W. (2021). AMRFinderPlus and the Reference Gene Catalog facilitate examination of the genomic links among antimicrobial resistance, stress response, and virulence. Scientific Reports, 11(1), 12728. 10.1038/s41598-021-91456-0

Foster, P. L., Lee, H., Popodi, E., Townes, J. P., & Tang, H. (2015). Determinants of spontaneous mutation in the bacterium *Escherichia coli* as revealed by whole-genome sequencing. Proceedings of the National Academy of Sciences, 112(44). 10.1073/pnas.1512136112

Froelich, J. M., Tran, K., & Wall, D. (2006). A pmrA constitutive mutant sensitizes Escherichia coli to deoxycholic acid. Journal of Bacteriology, 188(3), 1180–1183. 10.1128/JB.188.3.1180-1183.2006

Gifford, D. R., Berríos-Caro, E., Joerres, C., Suñé, M., Forsyth, J. H., Bhattacharyya, A., Galla, T., & Knight, C. G. (2023). Mutators can drive the evolution of multi-resistance to antibiotics. PLOS Genetics, 19(6), e1010791. 10.1371/journal.pgen.1010791

Gifford, D. R., Bhattacharyya, A., Geim, A., Marshall, E., Krašovec, R., & Knight, C. G. (2024). Environmental and genetic influence on the rate and spectrum of spontaneous mutations in Escherichia coli. Microbiology, 170(4). 10.1099/mic.0.001452

Hall, K. M., Williams, L. G., Smith, R. D., Kuang, E. A., Ernst, R. K., Bojanowski, C. M., Wimley, W. C., Morici, L. A., & Pursell, Z. F. (2025). Mutational signature analysis predicts bacterial hypermutation and multidrug resistance. Nature Communications, 16(1), 19. 10.1038/s41467-024-55206-w

Hall, L. M. C., & Henderson-Begg, S. K. (2006). Hypermutable bacteria isolated from humans - A critical analysis. In Microbiology (Vol. 152, Issue 9, pp. 2505–2514). 10.1099/mic.0.29079-0

Hughes, D., & Andersson, D. I. (2015). Evolutionary consequences of drug resistance: shared principles across diverse targets and organisms. Nature Reviews Genetics, 16(8), 459–471. 10.1038/nrg3922

Jin, D. J., & Gross, C. A. (1988). Mapping and sequencing of mutations in the Escherichia colirpoB gene that lead to rifampicin resistance. Journal of Molecular Biology, 202(1), 45–58. 10.1016/0022-2836(88)90517-7

Jolivet-Gougeon, A., Kovacs, B., le Gall-David, S., le Bars, H., Bousarghin, L., Bonnaure-Mallet, M., Lobel, B., Guillé, F., Soussy, C.-J., & Tenke, P. (2011). Bacterial hypermutation: clinical implications. Journal of Medical Microbiology, 60(5), 563–573. 10.1099/jmm.0.024083-0

Komp Lindgren, P., Karlsson, A., & Hughes, D. (2003). Mutation Rate and Evolution of Fluoroquinolone Resistance in *Escherichia coli* Isolates from Patients with Urinary Tract Infections. Antimicrobial Agents and Chemotherapy, 47(10), 3222–3232. 10.1128/AAC.47.10.3222-3232.2003

LeClerc, J. E., Li, B., Payne, W. L., & Cebula, T. A. (1996). High Mutation Frequencies Among *Escherichia coli* and *Salmonella* Pathogens. Science, 274(5290), 1208–1211. 10.1126/science.274.5290.1208

Leighow, S. M., Liu, C., Inam, H., Zhao, B., & Pritchard, J. R. (2020). Multi-scale Predictions of Drug Resistance Epidemiology Identify Design Principles for Rational Drug Design. Cell Reports, 30(12), 3951–3963.e4. 10.1016/j.celrep.2020.02.108

MacLean, R. C., Hall, A. R., Perron, G. G., & Buckling, A. (2010). The population genetics of antibiotic resistance: integrating molecular mechanisms and treatment contexts. Nature Reviews Genetics, 11(6), 405–414. 10.1038/nrg2778

Maharjan, R. P., & Ferenci, T. (2017). A shifting mutational landscape in 6 nutritional states: Stress-induced mutagenesis as a series of distinct stress input–mutation output relationships. PLOS Biology, 15(6), e2001477. 10.1371/journal.pbio.2001477

Martinez, J. L., & Baquero, F. (2000). Mutation Frequencies and Antibiotic Resistance. Antimicrobial Agents and Chemotherapy, 44(7), 1771–1777. 10.1128/AAC.44.7.1771-1777.2000

Mehta, H. H., Prater, A. G., Beabout, K., Elworth, R. A. L., Karavis, M., Gibbons, H. S., & Shamoo, Y. (2019). The Essential Role of Hypermutation in Rapid Adaptation to Antibiotic Stress. Antimicrobial Agents and Chemotherapy, 63(7). 10.1128/AAC.00744-19

Miller, K., O’Neill Alexander John, & Chopra Ian. (2002). Response of Escherichia coli hypermutators to selection pressure with antimicrobial agents from different classes. Journal of Antimicrobial Chemotherapy, 49(6), 925–934. 10.1093/jac/dkf044

Murray, C. J. L., Ikuta, K. S., Sharara, F., Swetschinski, L., Robles Aguilar, G., Gray, A., Han, C., Bisignano, C., Rao, P., Wool, E., Johnson, S. C., Browne, A. J., Chipeta, M. G., Fell, F., Hackett, S., Haines-Woodhouse, G., Kashef Hamadani, B. H., Kumaran, E. A. P., McManigal, B.,…Naghavi, M. (2022). Global burden of bacterial antimicrobial resistance in 2019: a systematic analysis. The Lancet, 399(10325), 629–655. 10.1016/S0140-6736(21)02724-0

Oliver, A., Cantón, R., Campo, P., Baquero, F., & Blázquez, J. (2000). High Frequency of Hypermutable *Pseudomonas aeruginosa* in Cystic Fibrosis Lung Infection. Science, 288(5469), 1251–1253. 10.1126/science.288.5469.1251

Parveen, S., Madhwal, A., Ruchith, B., Sane, M., & Agashe, D. (2025). Mutation bias is a key predictor of adaptation rate. 10.1101/2025.09.22.677663

Payne, J. L., Menardo, F., Trauner, A., Borrell, S., Gygli, S. M., Loiseau, C., Gagneux, S., & Hall, A. R. (2019). Transition bias influences the evolution of antibiotic resistance in Mycobacterium tuberculosis. PLOS Biology, 17(5), e3000265. 10.1371/journal.pbio.3000265

Pelchovich, G., Schreiber, R., Zhuravlev, A., & Gophna, U. (2013a). The contribution of common rpsL mutations in Escherichia coli to sensitivity to ribosome targeting antibiotics. International Journal of Medical Microbiology, 303(8), 558–562. 10.1016/j.ijmm.2013.07.006

Pelchovich, G., Schreiber, R., Zhuravlev, A., & Gophna, U. (2013b). The contribution of common rpsL mutations in Escherichia coli to sensitivity to ribosome targeting antibiotics. International Journal of Medical Microbiology, 303(8), 558–562. 10.1016/j.ijmm.2013.07.006

Pourahmad Jaktaji, R., & Mohiti, E. (2010). Study of Mutations in the DNA gyrase gyrA Gene of Escherichia coli. Iranian Journal of Pharmaceutical Research: IJPR, 9(1), 43–48.

Quesada, A., Porrero, M. C., Téllez, S., Palomo, G., García, M., & Domínguez, L. (2015). Polymorphism of genes encoding PmrAB in colistin-resistant strains of Escherichia coli and Salmonella enterica isolated from poultry and swine. Journal of Antimicrobial Chemotherapy, 70(1), 71–74. 10.1093/jac/dku320

R Core Team. (2024). R: A Language and Environment for Statistical Computing. R Foundation for Statistical Computing.

Ruis, C., Weimann, A., Tonkin-Hill, G., Pandurangan, A. P., Matuszewska, M., Murray, G. G. R., Lévesque, R. C., Blundell, T. L., Floto, R. A., & Parkhill, J. (2023). Mutational spectra are associated with bacterial niche. Nature Communications, 14(1), 7091. 10.1038/s41467-023-42916-w

Sane, M., Diwan, G. D., Bhat, B. A., Wahl, L. M., & Agashe, D. (2023). Shifts in mutation spectra enhance access to beneficial mutations. Proceedings of the National Academy of Sciences, 120(22). 10.1073/pnas.2207355120

Sane, M., Parveen, S., & Agashe, D. (2025). Mutation bias alters the distribution of fitness effects of mutations. PLOS Biology, 23(7), e3003282. 10.1371/journal.pbio.3003282

Shepherd, M. J., Fu, T., Harrington, N. E., Kottara, A., Cagney, K., Chalmers, J. D., Paterson, S., Fothergill, J. L., & Brockhurst, M. A. (2024). Ecological and evolutionary mechanisms driving within-patient emergence of antimicrobial resistance. Nature Reviews Microbiology, 22(10), 650–665. 10.1038/s41579-024-01041-1

Shewaramani, S., Finn, T. J., Leahy, S. C., Kassen, R., Rainey, P. B., & Moon, C. D. (2017). Anaerobically Grown Escherichia coli Has an Enhanced Mutation Rate and Distinct Mutational Spectra. PLOS Genetics, 13(1), e1006570. 10.1371/journal.pgen.1006570

Shibai, A., Izutsu, M., Kotani, H., & Furusawa, C. (2025). Quantitative analysis of relationship between mutation rate and speed of adaptation under antibiotic exposure in Escherichia coli. PLOS Genetics, 21(3), e1011627. 10.1371/journal.pgen.1011627

Sniegowski, P. D., Gerrish, P. J., & Lenski, R. E. (1997). Evolution of high mutation rates in experimental populations of E. coli. Nature, 387(6634), 703–705. 10.1038/42701

Stoltzfus, A., & McCandlish, D. M. (2017). Mutational Biases Influence Parallel Adaptation. Molecular Biology and Evolution, 34(9), 2163–2172. 10.1093/molbev/msx180

Tuffaha, M. Z., Castellano, D., Colomé, C. S., Gutenkunst, R. N., & Wahl, L. M. (2025). Nonhypermutator Cancers Access Driver Mutations Through Reversals in Germline Mutational Bias. Molecular Biology and Evolution, 42(5). 10.1093/molbev/msaf105

Tuffaha, M. Z., Varakunan, S., Castellano, D., Gutenkunst, R. N., & Wahl, L. M. (2023). Shifts in Mutation Bias Promote Mutators by Altering the Distribution of Fitness Effects. The American Naturalist, 202(4), 503–518. 10.1086/726010

van der Putten, B. C. L., Remondini, D., Pasquini, G., Janes, V. A., Matamoros, S., & Schultsz, C. (2019). Quantifying the contribution of four resistance mechanisms to ciprofloxacin MIC in *Escherichia coli*: a systematic review. Journal of Antimicrobial Chemotherapy, 74(2), 298–310. 10.1093/jac/dky417

Watanabe, R., & Doukyu, N. (2012). Contributions of mutations in acrR and marR genes to organic solvent tolerance in Escherichia coli. AMB Express, 2(1), 58. 10.1186/2191-0855-2-58

Wiegand, I., Hilpert, K., & Hancock, R. E. W. (2008). Agar and broth dilution methods to determine the minimal inhibitory concentration (MIC) of antimicrobial substances. Nature Protocols, 3(2), 163–175. 10.1038/nprot.2007.521

Woodford, N., & Ellington, M. J. (2007). The emergence of antibiotic resistance by mutation. Clinical Microbiology and Infection, 13(1), 5–18. 10.1111/j.1469-0691.2006.01492.x

