## Supplemental_file for "Mutation bias influences the emergence and effects of antibiotic resistance"

### SUPPLEMENTARY FIGURES

**Fig. S1. Cell density at a standardized optical density across strains.** Plots showing colony forming units (CFU) per mL measured in LB when ancestral cultures reached an $OD_{600}$ of 0.5 for each strain (mean ± SD, n = 3 biological replicates). One-way ANOVA detected no significant differences across strains ($F = 0.593$, $df = 6$, $p=0.731$), indicating that all strains achieved comparable cell densities at $OD_{600} = 0.5$.

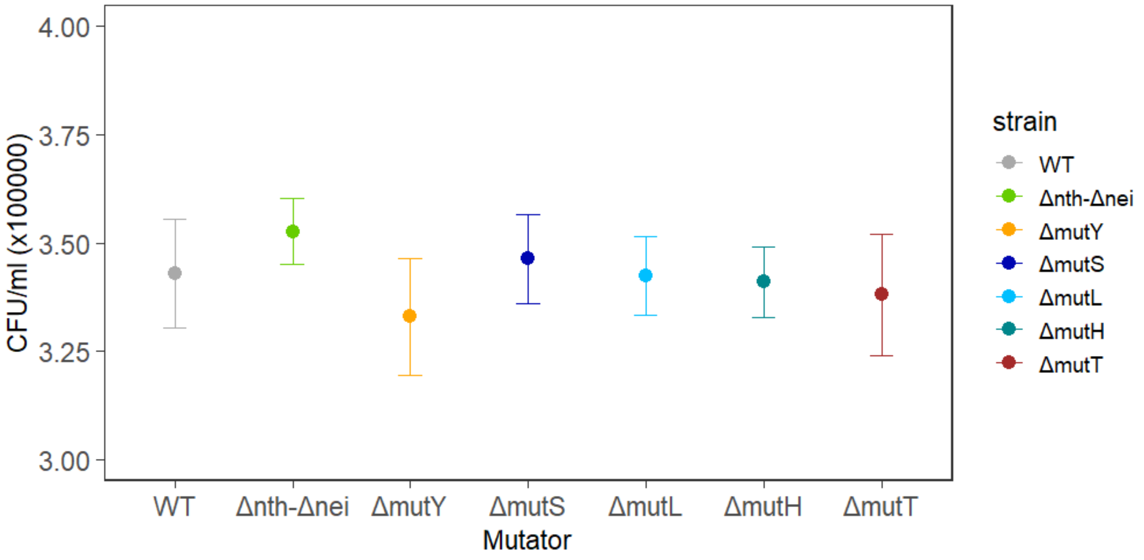

**Fig. S2. Growth curves for WT and mutator strains under antibiotic stress (Experiment 1).** Mean (± SD, n=3 biological replicates) of $OD_{600}$ over time for strains exposed to increasing concentrations of each antibiotic. Colours represent antibiotic concentrations, indicated in keys on the right.

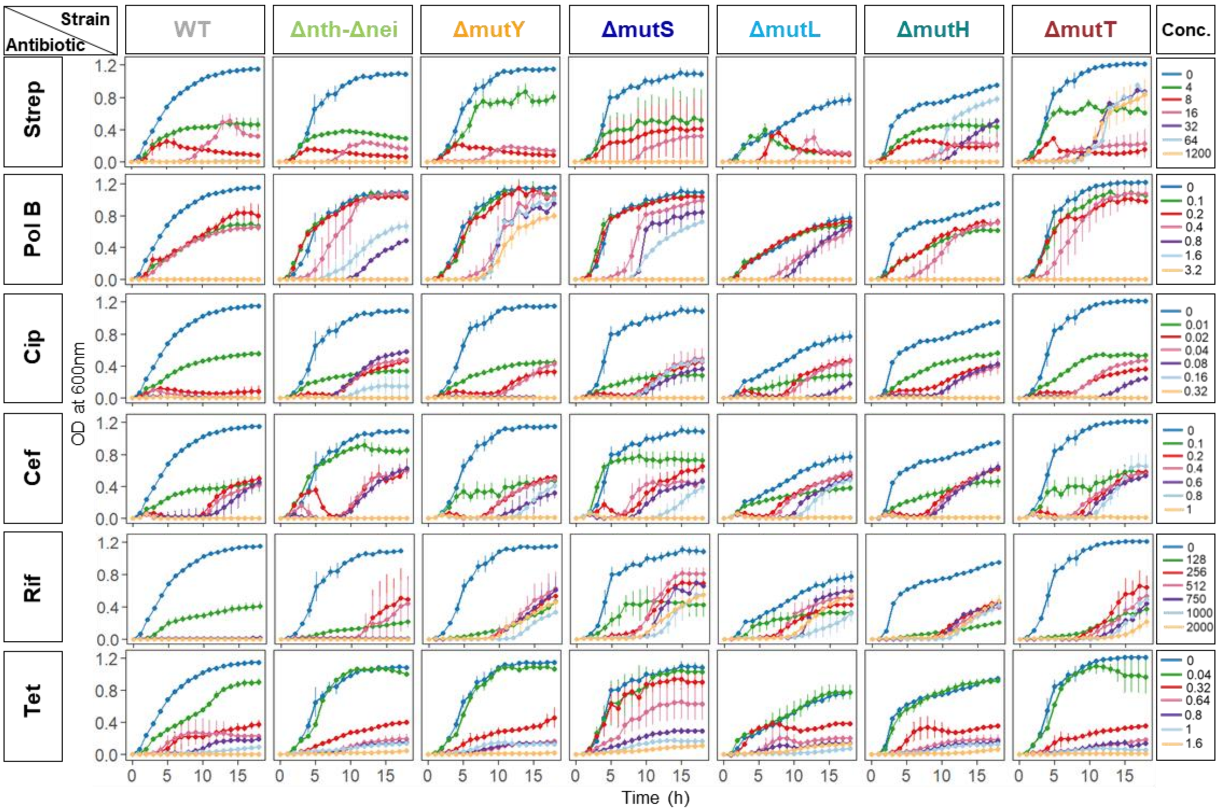

**Fig. S3: Testing for residual antibiotic in spent media.** Strains were cultured for 18 hours at the respective highest sub-MIC antibiotic concentrations used for Experiment 2, or in LB without antibiotic (control). These cultures were used to prepare spent media. From stocks, resistant populations were inoculated into spent LB either without antibiotic (n=6, black) or with antibiotic (n=3, maroon). The change in $OD_{600}$ values over time from this growth cycle is shown. Lines represent mean $OD_{600}$ across biological replicates and shaded regions indicate SD. Overall higher OD values of black lines compared to maroon lines indicate presence of residual antibiotic in the spent medium. Empty panels represent cases no resistance did not emerge during the first 18 hours of the experiment.

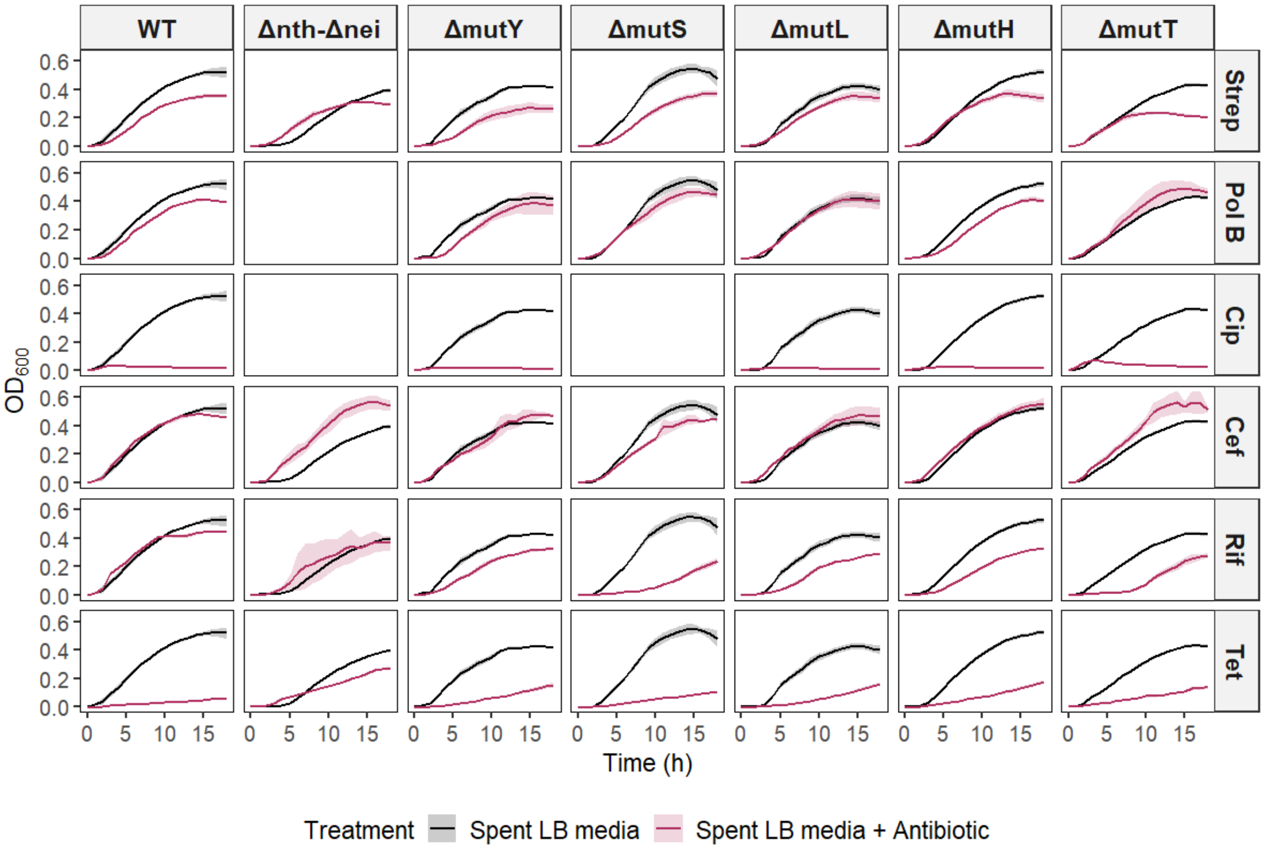

**Fig. S4. Optical density of cultures across antibiotic concentration gradients (Experiment 1).** Plots show the change in final $OD_{600}$ values of strains as a function of increasing concentrations of antibiotics (mean ± SD, n = 3 biological replicates). Colours denote strains.

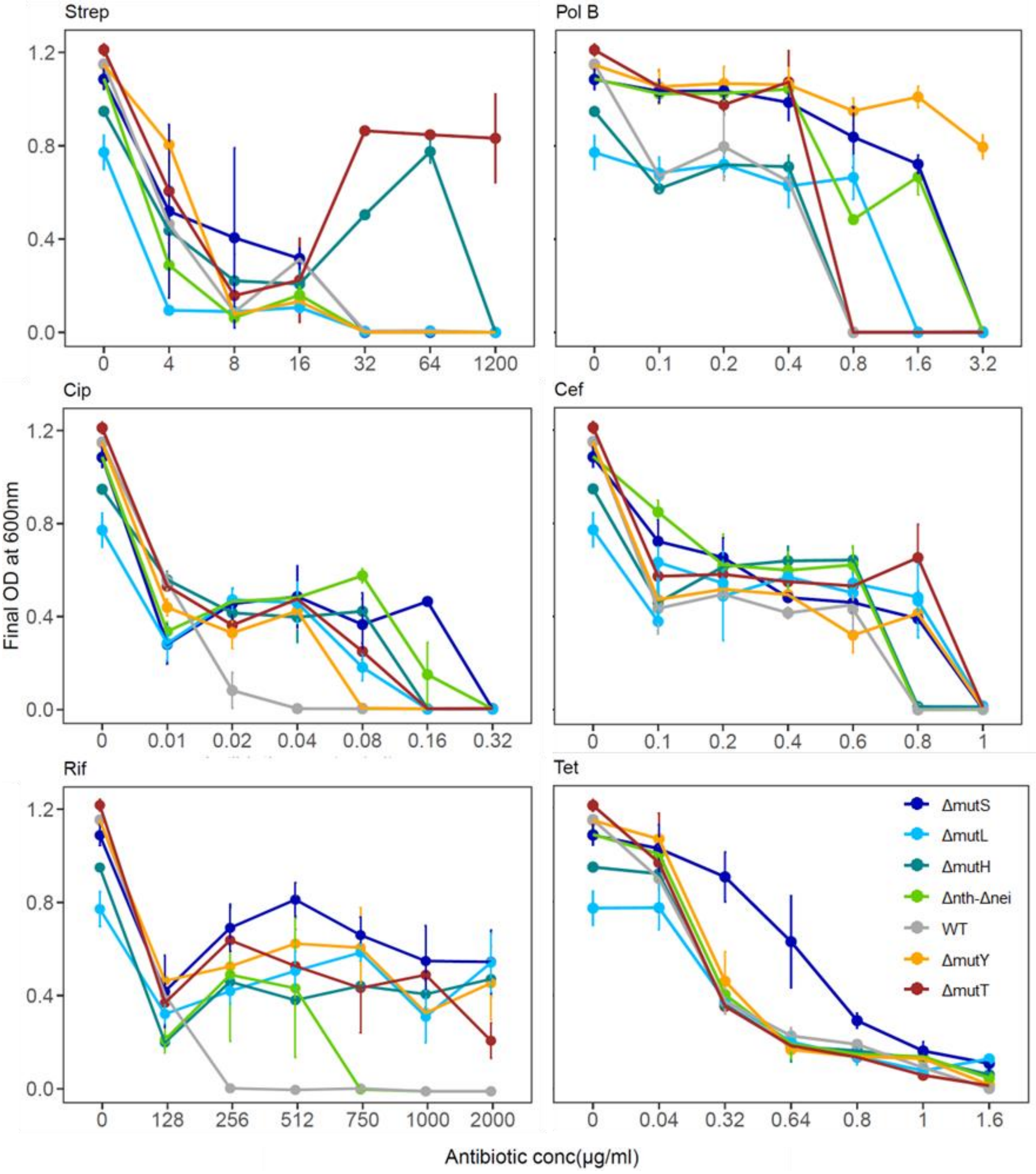

**Fig. S5. Growth rate of cultures across antibiotic concentration gradients (Experiment 1).** Plots show the change in growth rate values of strains as a function of increasing concentrations of antibiotics (mean ± SD, n = 3 biological replicates). Colours denote strains.

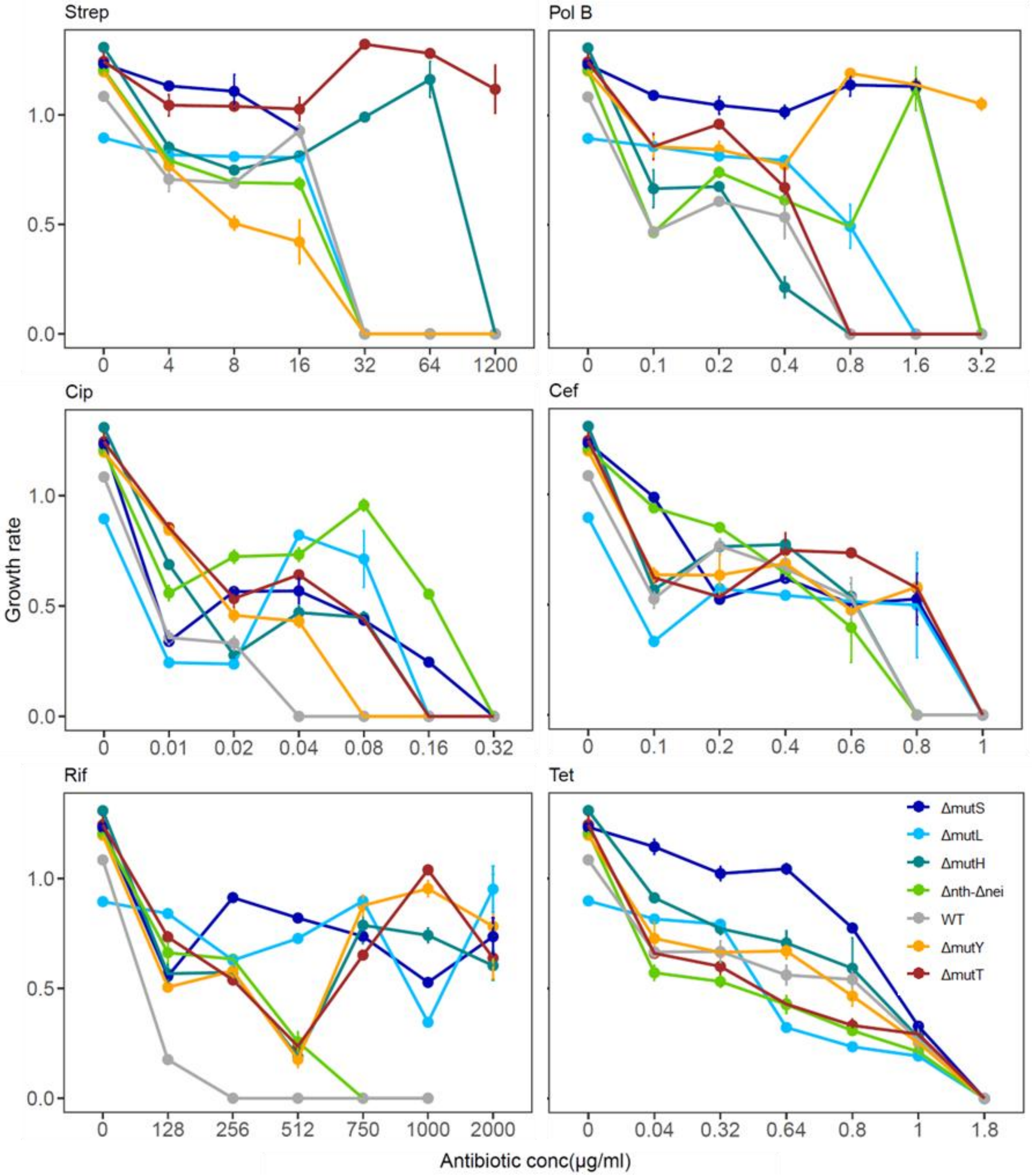

**Fig S6: Cost of antibiotic resistance mutations in absence of antibiotic (Experiment 3).** Cost was calculated as the difference between the final $OD_{600}$ of resistant populations (grown under the highest sub-MIC concentration (identified in Experiment 2) and that of their corresponding ancestral strains grown in antibiotic-free LB. Points represent differences in $OD_{600}$ between resistant mutants and respective ancestors (n=2-6, mean ± standard deviation). In two cases, only one replicate showed measurable growth (i.e., showed resistance); these are marked, and were not analysed statistically. The dashed horizontal line denotes no difference from the ancestor (i.e., no cost). Negative values indicate that resistance mutations reduced $OD_{600}$ relative to the ancestor (fitness cost), whereas positive values indicate a benefit in the absence of antibiotic. Asterisks indicate strain-antibiotic combinations for which costs were significantly different from zero (t-tests, $p < 0.05$).

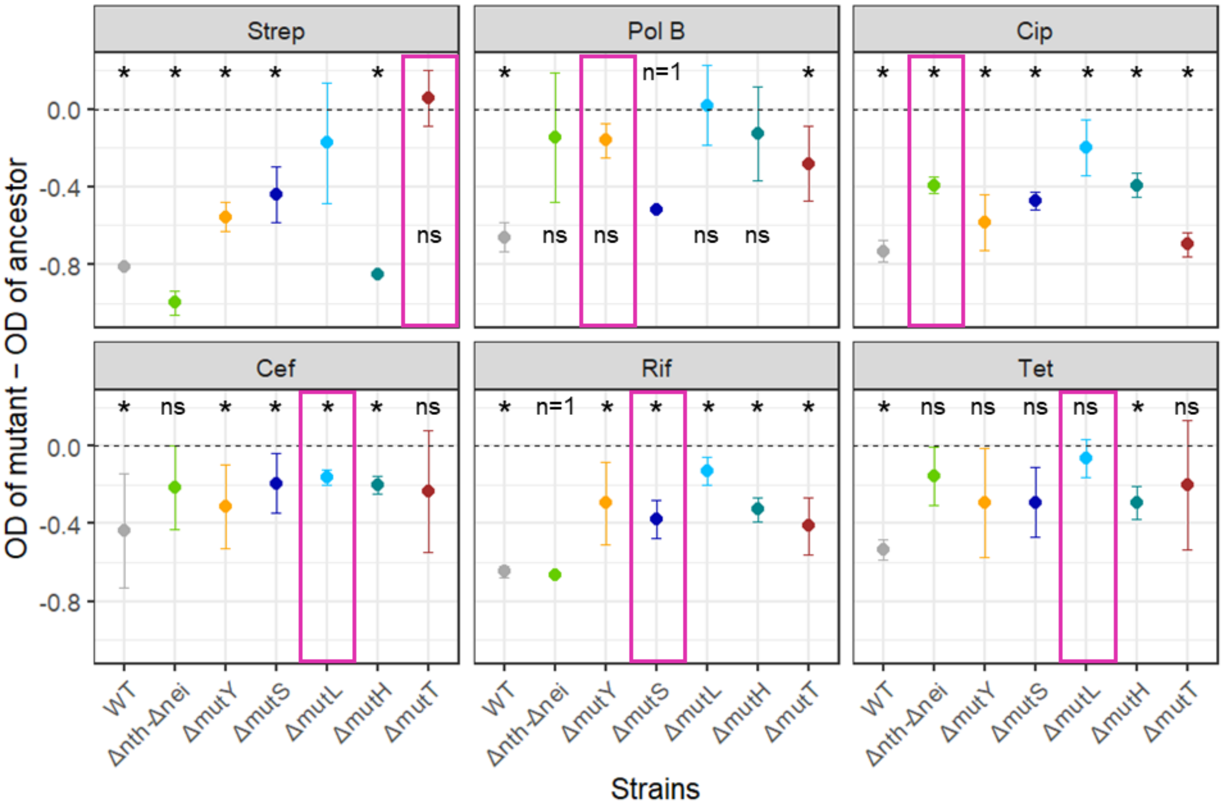


ns

**Fig. S7. Growth curves of replicate resistant populations at their highest sub-lethal antibiotic concentration.** Mean (± SD, n=3 biological replicates) of $OD_{600}$ over time for strains exposed to antibiotics at the concentrations used for Experiment 2. The number of replicates for each strain antibiotic concentration is variable (Table 2)

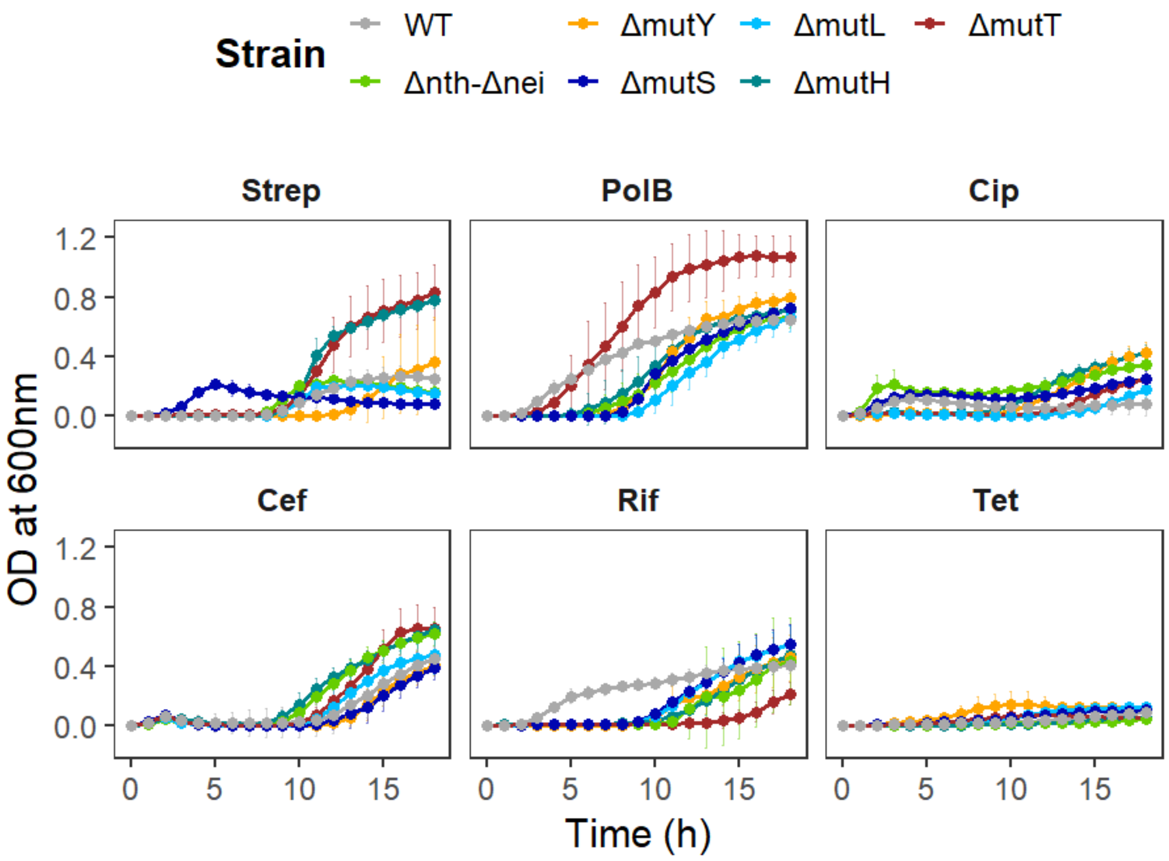

**Fig. S8: Mutations identified (using Sanger sequencing) in resistant populations for each antibiotic within target genes.** At the top, the mutation spectrum of each strain is indicated (based on mutation accumulation studies, MA); this is identical to Fig. 1D. For each antibiotic (indicated on the far right), the occurrence of mutations in target genes is shown, for each of six replicate populations per strain with sanger sequencing (represented in columns). Each row indicates a distinct mutation in the gene indicated on the left. Repeated gene names for a given strain and antibiotic combination indicate mutations observed at different positions in the same gene. However, across strains, the row identity is not meaningful. E.g., in Rif, several strains have *rpoB* mutations (first row); these are not necessarily at the same position across strains. No relevant target genes are known for Tet, and for Cef, we did not find any mutations in the predicted target genes.

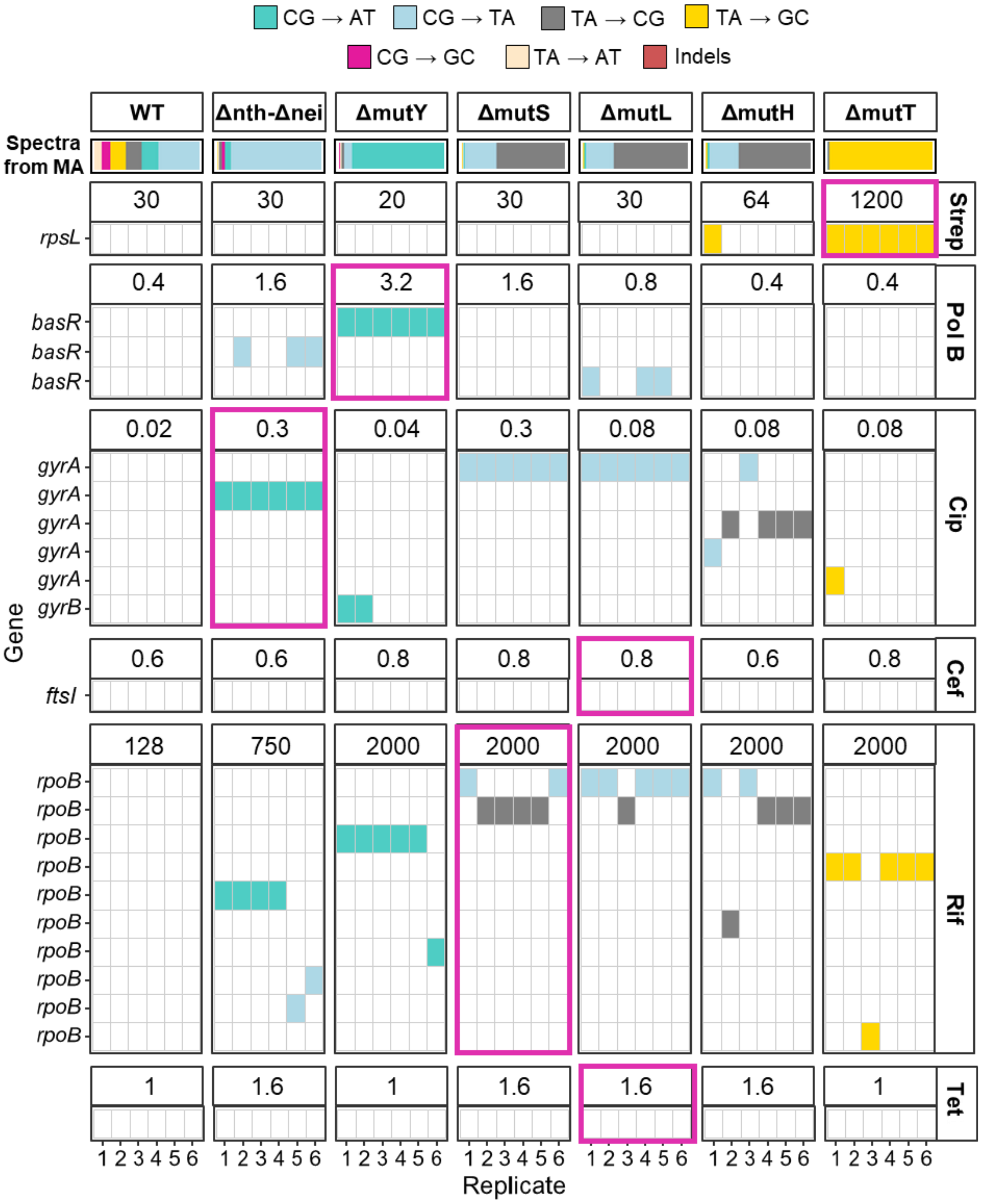

**Fig. S9: The spectrum of efflux pump related mutations observed in resistant populations of mutators in each antibiotic.** At the top, the mutation spectrum of each strain is indicated (based on mutation accumulation studies, MA); this is identical to Fig. 1D. For each antibiotic (indicated on the far right), the occurrence of mutations in known efflux pump related genes is shown, for each of three replicate populations per strain with NGS data (represented in columns). Numbers above each panel indicate the highest sub-MIC concentration (ug/mL) at which the strain was tested (Experiment 2). Each row indicates a distinct mutation in the gene indicated on the left. Repeated gene names for a given strain and antibiotic combination indicate mutations observed at different positions in the same gene. However, across strains, the row identity is not meaningful. E.g., in Cef, several strains have *marR* mutations (first row); these are not necessarily at the same position across strains. Numbers inside each cell indicate the frequency at which the mutation was observed (a value of 1 indicates that the allele was fixed in the population). Pink boxes around panels indicate the fittest mutator in each antibiotic.

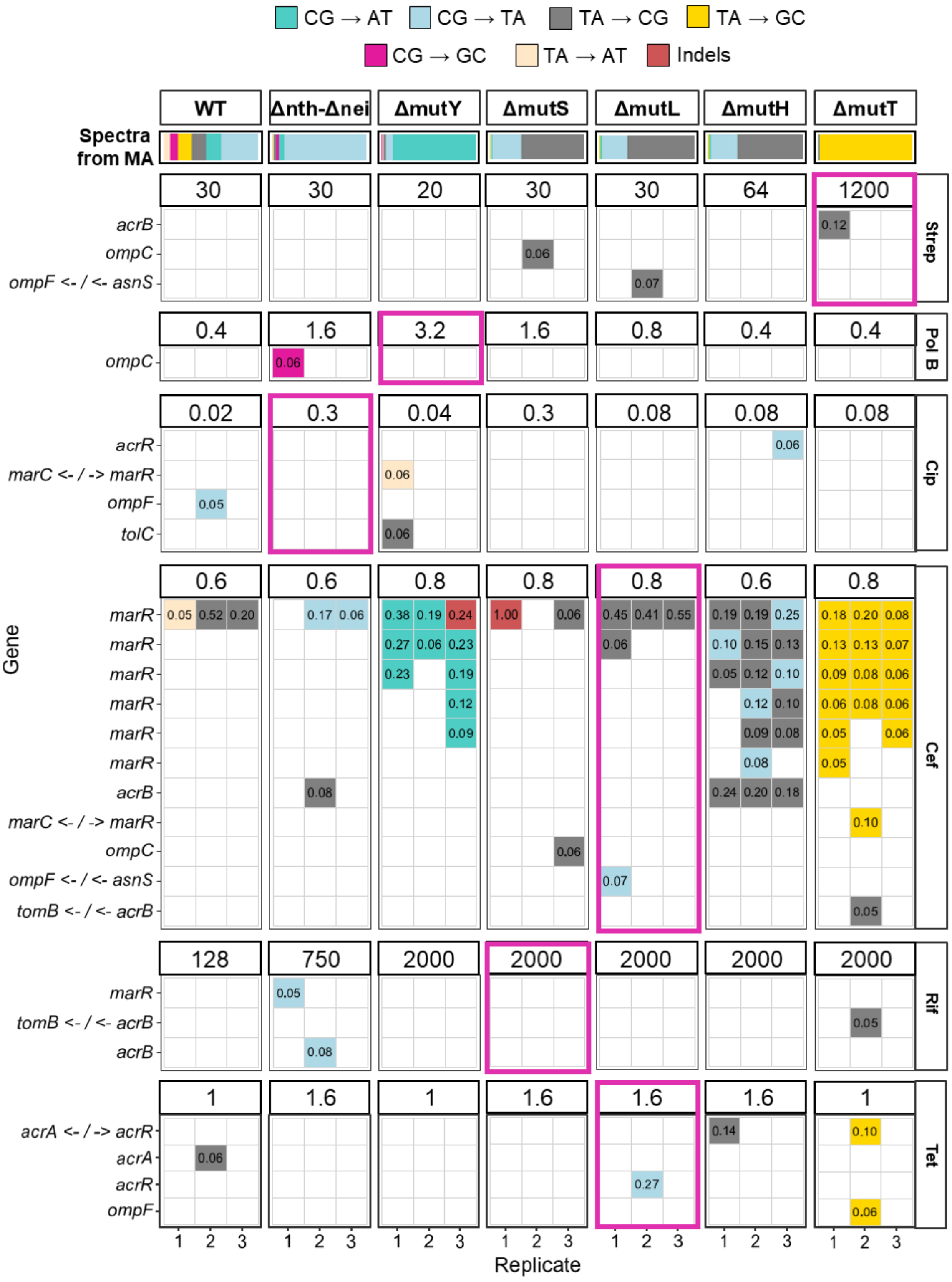

CG → AT
CG → TA
TA → CG
TA → GC
CG → GC
TA → AT
Indels
WT
Δnth-Δnei
ΔmutY
ΔmutS
ΔmutL
ΔmutH
ΔmutT
Spectra from MA
Strep
Pol B
Cip
Cef
Rif
Tet
Gene
Replicate
acrB
ompC
ompF <- / <- asnS
marC <- / -> marR
acrR
ompF
tolC
marR
tomB <- / <- acrB
acrA <- / -> acrR
acrA

**Fig. S10: The spectrum of mutations observed in target genes of other antibiotics in resistant populations of mutators in each antibiotic.** At the top, the mutation spectrum of each strain is indicated; this is identical to Fig. 1D. For each antibiotic (indicated on the far right), the occurrence of mutations in known target genes of other antibiotics is shown, for each of three replicate populations per strain with NGS data (represented in columns). Numbers above each panel indicate the highest sub-MIC concentration (μg/mL) at which the strain was tested (Experiment 2). Each row indicates a distinct mutation in the gene indicated on the left. Numbers inside each cell indicate the frequency at which the mutation was observed (a value of 1 indicates that the allele was fixed in the population). Pink boxes around panels indicate the fittest mutator in each antibiotic.

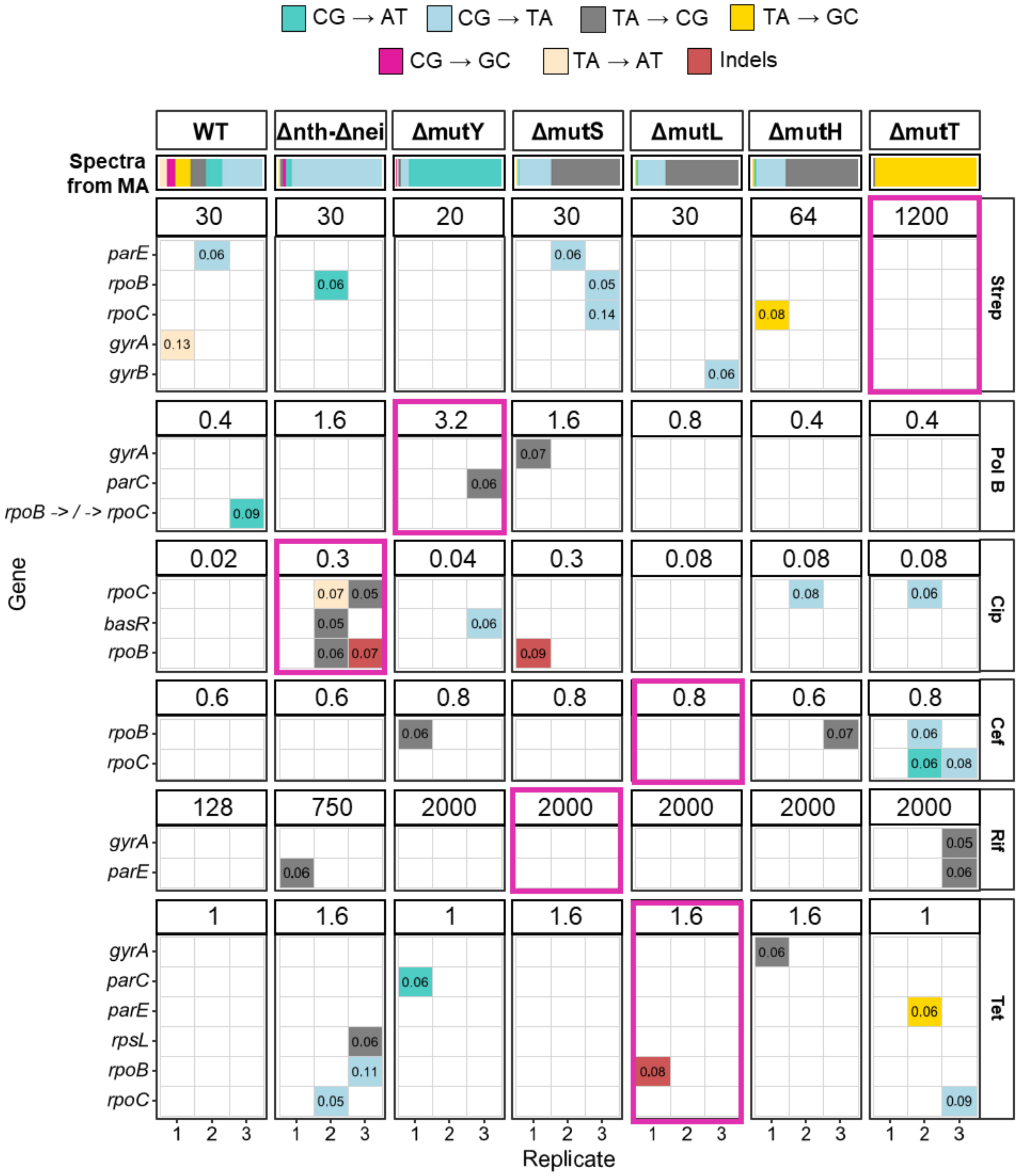

**Fig. S11: Mutation spectra of all strains according to mutation accumulation** (Sane et al., 2025). **(A)** Mutation spectra of each strain depicting proportions of different classes of single nucleotide substitutions (same as Fig. 1D) but including indels (not shown in Fig. 1D) **(B)** Mutation spectra of each strain depicting proportions of single nucleotide substitutions and indels. Total number of mutations for each strain in mentioned under the stacked bars.

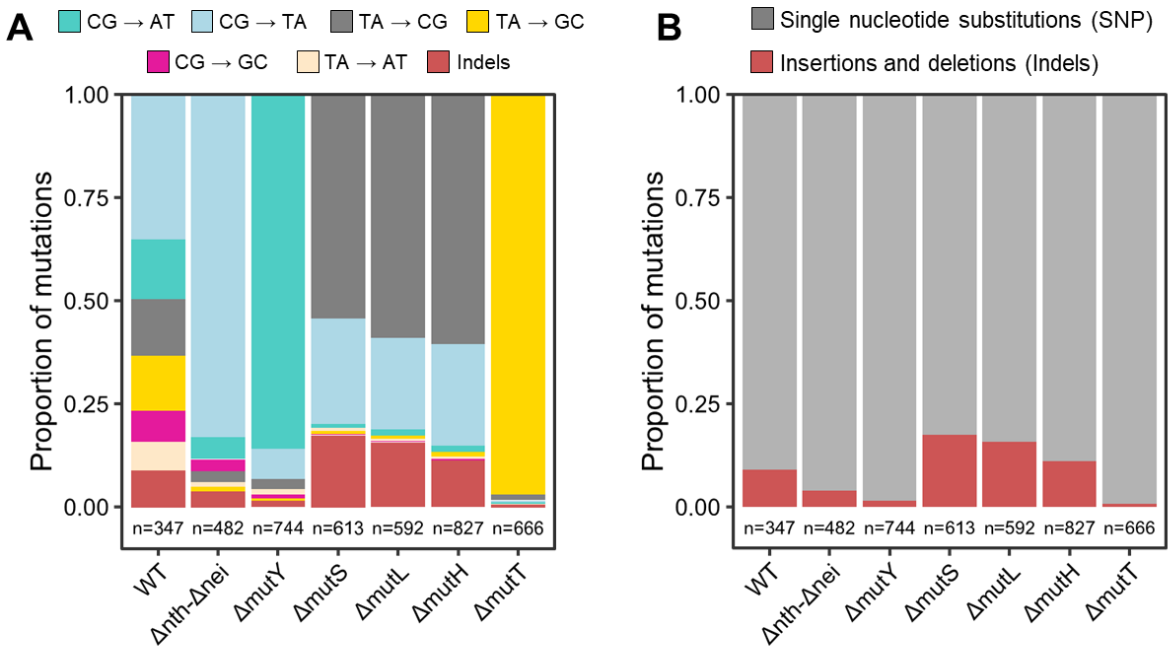

**Fig. S12: Mutational spectra of resistant populations across mutator backgrounds and antibiotics consisting of mutations above 80% frequency.** Stacked bars show the proportion of mutations belonging to different mutational classes in resistant populations of strains evolved under exposure to antibiotics. Each bar represents an independent replicate population (n=3), with colours denoting mutation classes. Numbers within bars indicate the total number of mutations detected in each population. Columns on the far right indicate mutation proportions observed from mutation accumulation experiments.

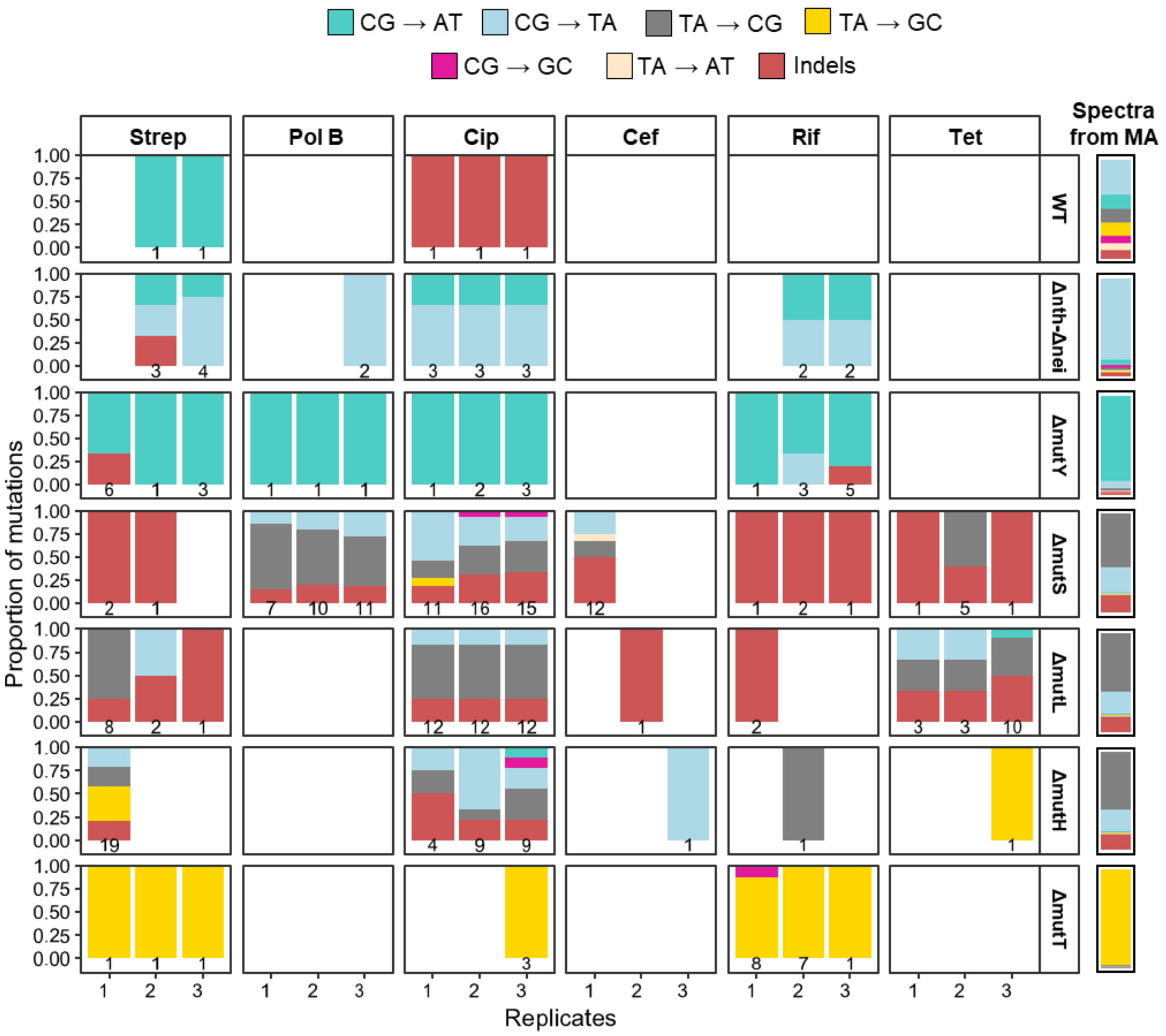

**Fig. S13: Mutational spectra of resistant populations across mutator backgrounds and antibiotics.** Stacked bars (n=3) show the proportion of mutations belonging to different mutational classes in resistant populations of strains evolved under exposure to highest sub-MIC antibiotic concentrations. Numbers within bars indicate the total number of mutations detected in each population. Columns on the far right indicate mutation proportions observed from mutation accumulation experiments.

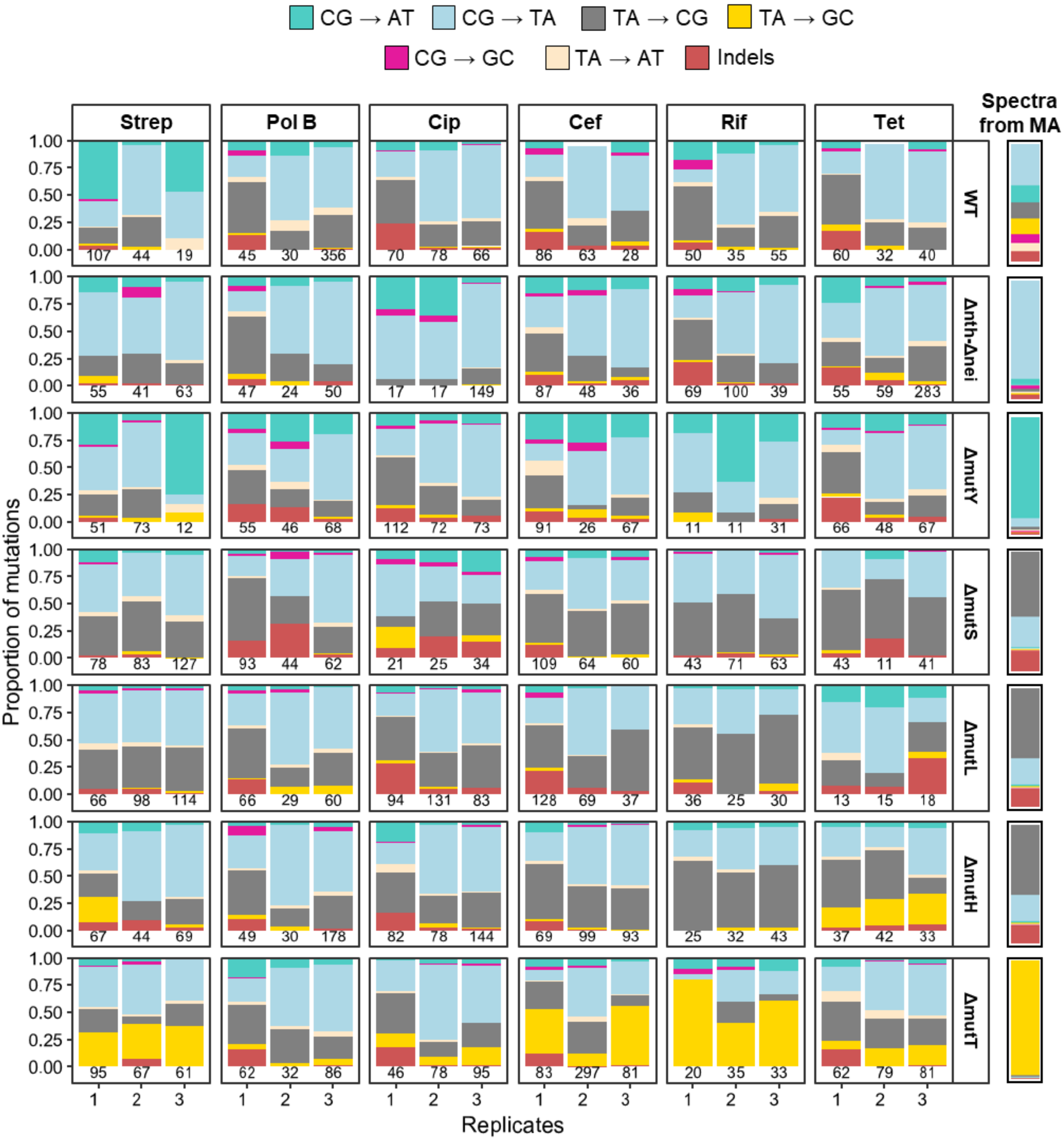

### SUPPLEMENTARY TABLES

**Table S1:** PCR primers used for amplification of antibiotic resistance-associated genes, as well as efflux pump genes.

| Antibiotic | Target gene | Forward primer (5'→3') | Reverse primer (5'→3') | Amplicon size | Amplification temperature (ºC) | Reference |
|---|---|---|---|---|---|---|
| **Strep** | *rpsL* | ATGGCAACAGTTAACCAGCT | CCTTAGGACGCTTCACGC | 375 | 60 | (Pelchovich et. al, 2013) |
| **PolB** | *basR* | AGTTTTCCTCATTCGCGACCA | TACCAGGCTGCGGATGATATTCT | 714 | 55 | (Quesada et. al, 2015) |
| **Cip** | *gyrA* | CTGAAGCCGGTACACCGT | GGATATACACCTTGCCGC | 590 | 56 | (Jaktaji and Mohiti, 2010) |
| **Cef** | *ftsI* | GCCCAGCATGTTTCACAAGATG | CGAGCAGAGATGCTGCGAA | 1704 | 60 | This study |
| **Rif** | *rpoB* | CGTGCGGTGAAAGAGCGTCTGTCT | ACG TTTAGCTACCGCAGTTACACC | 744 | 62 | This study |
| **Efflux** | *acrA* | CCAATTTGAAATCGGACACTCG | GCATGTCTTAACGGCTCCTG | 1187 | 60 | (Watanabe and Doukyu, 2012) |
| **Efflux** | *acrB* | GAAAGGCCAACAGCTTAAC | GAGCTGGAGTCAGGATCAAC | 800 | 55 | (Chowdhury et. al, 2019) |
| **Efflux** | *marR* | CTGTTCATGTTGCCTGCCAG | CAGTCCAAAATGCTATGAATGG | 825 | 52 | (Watanabe and Doukyu, 2012) |

**Table S2:** Mutations identified in target genes at intermediate and low antibiotic concentrations using Sanger sequencing, for a subset of strains and antibiotics. One colony each from 3 biological replicate populations was used for sequencing.

| Antibiotic | Target gene | Mutator | Concentration (μg/ml) | Mutation | Position in gene | Frequency of mutations (n=3) |
|---|---|---|---|---|---|---|
| Streptomycin | *rpsL* | ΔmutT | 64 | A → C | 129 | 1 |
| | | ΔmutT | 4 | A → C | 129 | 0.3 |
| Rifamycin | *rpoB* | ΔmutT | 1000 | T→ G | 1576 | 1 |
| | | ΔmutY | 1000 | G → T | 1546 | 1 |
| | | ΔmutT | 128 | A → C | 1547 | 0.3 |
| | | ΔmutY | 128 | C → A | 1537 | 0.3 |
| | | | 128 | C → A | 1576 | 0.3 |
| Ciprofloxacin | *gyrA* | Δnth-Δnei | 0.02 | --- | --- | 0 |
| Polymyxin B | *basR* | ΔmutY | 1.6 | C → A | 157 | 1 |
| | | | 0.2 | --- | --- | 0 |

**Table S3:** Sequencing depth of samples after whole genome sequencing, estimated from mapped reads obtained using breseq v.0.39.0 (Deatherage and Barrick, 2014) with default parameters.

| Antibiotic | Strain | Replicate | Total number of reads | Sequencing depth(X) |
|---|---|---|---|---|
| Strep | WT | 1 | 4999108 | 107.7011 |
| Strep | WT | 2 | 5310754 | 114.4152 |
| Strep | WT | 3 | 7917316 | 170.5711 |
| Strep | Δnth-Δnei | 1 | 5079309 | 109.4289 |
| Strep | Δnth-Δnei | 2 | 7803767 | 168.1248 |
| Strep | Δnth-Δnei | 3 | 7462600 | 160.7747 |
| Strep | ΔmutY | 1 | 5029289 | 108.3513 |
| Strep | ΔmutY | 2 | 7056796 | 152.032 |
| Strep | ΔmutY | 3 | 5108915 | 110.0667 |
| Strep | ΔmutS | 1 | 4866481 | 104.8437 |
| Strep | ΔmutS | 2 | 7735523 | 166.6545 |
| Strep | ΔmutS | 3 | 6234669 | 134.32 |
| Strep | ΔmutL | 1 | 5275386 | 113.6532 |
| Strep | ΔmutL | 2 | 5912726 | 127.3841 |
| Strep | ΔmutL | 3 | 7259830 | 156.4062 |
| Strep | ΔmutH | 1 | 5353772 | 115.342 |
| Strep | ΔmutH | 2 | 8019321 | 172.7687 |
| Strep | ΔmutH | 3 | 6895352 | 148.5538 |
| Strep | ΔmutT | 1 | 5220398 | 112.4685 |
| Strep | ΔmutT | 2 | 7600291 | 163.7411 |
| Strep | ΔmutT | 3 | 7706100 | 166.0206 |
| Pol B | WT | 1 | 6803055 | 146.5654 |
| Pol B | WT | 2 | 5815024 | 125.2792 |
| Pol B | WT | 3 | 4318247 | 93.03255 |
| Pol B | Δnth-Δnei | 1 | 6237127 | 134.373 |
| Pol B | Δnth-Δnei | 2 | 7097869 | 152.9169 |
| Pol B | Δnth-Δnei | 3 | 7581781 | 163.3423 |
| Pol B | ΔmutY | 1 | 6067283 | 130.7139 |
| Pol B | ΔmutY | 2 | 6856567 | 147.7182 |
| Pol B | ΔmutY | 3 | 8379125 | 180.5203 |
| Pol B | ΔmutS | 1 | 6992998 | 150.6575 |
| Pol B | ΔmutS | 2 | 6507283 | 140.1933 |
| Pol B | ΔmutS | 3 | 7361735 | 158.6016 |
| Pol B | ΔmutL | 1 | 7539354 | 162.4282 |
| Pol B | ΔmutL | 2 | 6133952 | 132.1502 |
| Pol B | ΔmutL | 3 | 7857230 | 169.2766 |
| Pol B | ΔmutH | 1 | 5757425 | 124.0383 |
| Pol B | ΔmutH | 2 | 6725831 | 144.9017 |
| Pol B | ΔmutH | 3 | 4727959 | 101.8594 |
| Pol B | ΔmutT | 1 | 5765161 | 124.2049 |
| Pol B | ΔmutT | 2 | 6101354 | 131.4479 |
| Pol B | ΔmutT | 3 | 3757031 | 80.94168 |

**Table S3:** *continued..*

| Antibiotic | Strain | Replicate | Total number of reads | Sequencing depth(X) |
|---|---|---|---|---|
| Cip | WT | 1 | 6420699 | 138.3279 |
| Cip | WT | 2 | 7180089 | 154.6882 |
| Cip | WT | 3 | 8642500 | 186.1945 |
| Cip | Δnth-Δnei | 1 | 5040259 | 162.8814 |
| Cip | Δnth-Δnei | 2 | 5564107 | 179.8101 |
| Cip | Δnth-Δnei | 3 | 2614055 | 84.47601 |
| Cip | ΔmutY | 1 | 5548552 | 119.5383 |
| Cip | ΔmutY | 2 | 7849131 | 169.1021 |
| Cip | ΔmutY | 3 | 7586404 | 163.4419 |
| Cip | ΔmutS | 1 | 6433735 | 207.9131 |
| Cip | ΔmutS | 2 | 6144159 | 198.5551 |
| Cip | ΔmutS | 3 | 5321568 | 171.9722 |
| Cip | ΔmutL | 1 | 7645046 | 164.7053 |
| Cip | ΔmutL | 2 | 6684982 | 144.0216 |
| Cip | ΔmutL | 3 | 7289484 | 157.045 |
| Cip | ΔmutH | 1 | 6576637 | 141.6874 |
| Cip | ΔmutH | 2 | 7601271 | 163.7622 |
| Cip | ΔmutH | 3 | 6741615 | 145.2417 |
| Cip | ΔmutT | 1 | 7067735 | 152.2677 |
| Cip | ΔmutT | 2 | 7907546 | 170.3606 |
| Cip | ΔmutT | 3 | 8260771 | 177.9705 |
| Cef | WT | 1 | 6123838 | 131.9323 |
| Cef | WT | 2 | 6823814 | 147.0126 |
| Cef | WT | 3 | 6653682 | 143.3473 |
| Cef | Δnth-Δnei | 1 | 6468723 | 139.3625 |
| Cef | Δnth-Δnei | 2 | 7609716 | 163.9441 |
| Cef | Δnth-Δnei | 3 | 6570161 | 141.5479 |
| Cef | ΔmutY | 1 | 5929505 | 127.7456 |
| Cef | ΔmutY | 2 | 8687289 | 187.1594 |
| Cef | ΔmutY | 3 | 6707412 | 144.5048 |
| Cef | ΔmutS | 1 | 5577714 | 120.1666 |
| Cef | ΔmutS | 2 | 8698104 | 187.3924 |
| Cef | ΔmutS | 3 | 5604176 | 120.7367 |
| Cef | ΔmutL | 1 | 5776469 | 124.4486 |
| Cef | ΔmutL | 2 | 7247368 | 156.1377 |
| Cef | ΔmutL | 3 | 7532707 | 162.285 |
| Cef | ΔmutH | 1 | 8398833 | 180.9449 |
| Cef | ΔmutH | 2 | 7768379 | 167.3624 |
| Cef | ΔmutH | 3 | 7951072 | 171.2983 |
| Cef | ΔmutT | 1 | 7673669 | 165.3219 |
| Cef | ΔmutT | 2 | 3735229 | 80.47197 |
| Cef | ΔmutT | 3 | 7858521 | 169.3044 |

**Table S3:** *continued..*

| Antibiotic | Strain | Replicate | Total number of reads | Sequencing depth(X) |
|---|---|---|---|---|
| Rif | WT | 1 | 6293976 | 135.5978 |
| Rif | WT | 2 | 8708146 | 187.6088 |
| Rif | WT | 3 | 7652433 | 164.8644 |
| Rif | Δnth-Δnei | 1 | 6542572 | 140.9535 |
| Rif | Δnth-Δnei | 2 | 6771402 | 145.8834 |
| Rif | Δnth-Δnei | 3 | 9282687 | 199.9867 |
| Rif | ΔmutY | 1 | 5009676 | 161.8931 |
| Rif | ΔmutY | 2 | 6624745 | 214.0858 |
| Rif | ΔmutY | 3 | 3650977 | 117.9853 |
| Rif | ΔmutS | 1 | 5710856 | 184.5525 |
| Rif | ΔmutS | 2 | 5555464 | 179.5308 |
| Rif | ΔmutS | 3 | 6698773 | 216.4781 |
| Rif | ΔmutL | 1 | 5220092 | 168.6929 |
| Rif | ΔmutL | 2 | 5906309 | 190.8688 |
| Rif | ΔmutL | 3 | 6527212 | 210.9339 |
| Rif | ΔmutH | 1 | 6720619 | 217.1841 |
| Rif | ΔmutH | 2 | 6937079 | 224.1792 |
| Rif | ΔmutH | 3 | 6673311 | 215.6553 |
| Rif | ΔmutT | 1 | 5868035 | 189.6319 |
| Rif | ΔmutT | 2 | 6546386 | 211.5535 |
| Rif | ΔmutT | 3 | 5806711 | 187.6501 |
| Tet | WT | 1 | 6866466 | 147.9315 |
| Tet | WT | 2 | 7341368 | 158.1628 |
| Tet | WT | 3 | 7554366 | 162.7517 |
| Tet | Δnth-Δnei | 1 | 3132852 | 101.2415 |
| Tet | Δnth-Δnei | 2 | 5300469 | 171.2904 |
| Tet | Δnth-Δnei | 3 | 5171128 | 167.1106 |
| Tet | ΔmutY | 1 | 7591099 | 163.543 |
| Tet | ΔmutY | 2 | 8122564 | 174.993 |
| Tet | ΔmutY | 3 | 7030458 | 151.4646 |
| Tet | ΔmutS | 1 | 5002149 | 161.6499 |
| Tet | ΔmutS | 2 | 7486432 | 241.9321 |
| Tet | ΔmutS | 3 | 6138638 | 198.3767 |
| Tet | ΔmutL | 1 | 4990324 | 161.2677 |
| Tet | ΔmutL | 2 | 6339148 | 204.8564 |
| Tet | ΔmutL | 3 | 5668636 | 183.1881 |
| Tet | ΔmutH | 1 | 6215566 | 200.8627 |
| Tet | ΔmutH | 2 | 5217763 | 168.6176 |
| Tet | ΔmutH | 3 | 5890856 | 190.3694 |
| Tet | ΔmutT | 1 | 6324398 | 136.2532 |
| Tet | ΔmutT | 2 | 6170030 | 132.9275 |
| Tet | ΔmutT | 3 | 6585554 | 141.8795 |

**Table S4:** Statistical tests for results shown in Fig. 2B-E. **(A**) Results of Chi square tests to compare the probability of resistance emergence between strains growing in the same concentration of antibiotic (Fig. 2B) **(B)** Results of unpaired t-tests to compare lag time, final $OD_{600}$ and growth rate between strains growing in the same concentration of antibiotic (Fig. 2C). Bold p-values indicate significant differences ($p \leq 0.05$).

**A. Chi square test results for Fig. 2B**

| Antibiotic | Tested pair | Chi square statistic | df | p value |
|---|---|---|---|---|
| Strep | ΔmutT vs. ΔmutH | 75.908 | 1 | **2.20E-16** |
| Pol B | Δnth-Δnei vs. ΔmutS | 1.4945 | 1 | 0.2215 |
| Cip | Δnth-Δnei vs. ΔmutS | 0.26471 | 1 | 0.6069 |
| Cef | ΔmutY vs. ΔmutT | 6.451 | 1 | **0.01109** |
| | ΔmutY vs. ΔmutL | 21.385 | 1 | **3.76E-06** |
| | ΔmutY vs. ΔmutS | 35.767 | 1 | **2.22E-09** |
| | ΔmutT vs. ΔmutL | 4.2698 | 1 | **0.03879** |
| | ΔmutT vs. ΔmutS | 14.295 | 1 | **0.00016** |
| | ΔmutL vs. ΔmutS | 3.3882 | 1 | 0.06566 |
| Rif | ΔmutY vs. ΔmutT | 0 | 1 | 1 |
| | ΔmutY vs. ΔmutH | 6.9698 | 1 | **0.00829** |
| | ΔmutY vs. ΔmutL | 6.9698 | 1 | **0.00829** |
| | ΔmutY vs. ΔmutS | 6.9698 | 1 | **0.00829** |
| | ΔmutT vs. ΔmutH | 5.753 | 1 | **0.01646** |
| | ΔmutT vs. ΔmutL | 5.753 | 1 | **0.01646** |
| | ΔmutT vs. ΔmutS | 5.753 | 1 | **0.01646** |
| | ΔmutH vs. ΔmutL | NA | 1 | NA |
| | ΔmutH vs. ΔmutS | NA | 1 | NA |
| | ΔmutL vs. ΔmutS | NA | 1 | NA |
| Tet | Δnth-Δnei vs. ΔmutS | 4.6296 | 1 | **0.03142** |
| | Δnth-Δnei vs. ΔmutH | 4.6296 | 1 | **0.03142** |
| | Δnth-Δnei vs. ΔmutL | 4.6296 | 1 | **0.03142** |

**B. t-test results for Fig. 2C-E**

| Parameter | Antibiotic | Antibiotic concentration | Strain 1 | Strain 2 | p_value | t statistic |
|---|---|---|---|---|---|---|
| Lag time | Cip | 0.3 | Δnth-Δnei | ΔmutS | **1.42119E-05** | 5.237229366 |
| Lag time | Cef | 0.8 | ΔmutL | ΔmutS | **0.024057693** | -2.498891312 |
| Lag time | Rif | 2000 | ΔmutH | ΔmutL | 0.124718334 | 1.565714178 |
| Lag time | Rif | 2000 | ΔmutH | ΔmutS | **0.010855985** | 2.846836016 |
| Lag time | Rif | 2000 | ΔmutL | ΔmutS | **0.049532192** | 2.07129145 |
| Lag time | Tet | 1.6 | ΔmutH | ΔmutL | **1.55824E-15** | 12.04813963 |
| Lag time | Tet | 1.6 | ΔmutH | ΔmutS | **1.71368E-16** | 13.14055803 |
| Lag time | Tet | 1.6 | ΔmutL | ΔmutS | 0.161045744 | 1.485030253 |
| Final OD | Tet | 1.6 | ΔmutL | ΔmutS | **0.006046496** | 3.040353637 |
| Growth rate | Tet | 1.6 | ΔmutL | ΔmutS | **8.82E-10** | 8.611863581 |

**Table S5:** Target gene mutations from whole genome sequencing at intermediate antibiotic concentrations (Cip: 0.16 μg/ml, Rif: 1000 μg/ml and Tet: 1 μg/ml). Mutations were called using breseq v.0.39.0 (Deatherage and Barrick, 2014) with default parameters.

| Antibiotic | Strains | Replicate | Position | Mutation | Annotation | Gene | Frequency |
|---|---|---|---|---|---|---|---|
| cip | Δnth-Δnei | 1 | 2338460 | A→T | Y321N (TAC→AAC) | gyrA ← | 0.144 |
| cip | Δnth-Δnei | 1 | 2339173 | G→A | S83L (TCG→TTG) | gyrA ← | 1 |
| cip | Δnth-Δnei | 1 | 3878922 | A→G | L400L (TTA→CTA) | gyrB ← | 0.069 |
| cip | Δnth-Δnei | 2 | 2337574 | C→A | G616V (GGT→GTT) | gyrA ← | 0.115 |
| cip | Δnth-Δnei | 2 | 2339080 | C→A | G114V (GGC→GTC) | gyrA ← | 0.128 |
| cip | Δnth-Δnei | 2 | 2339173 | G→A | S83L (TCG→TTG) | gyrA ← | 1 |
| cip | Δnth-Δnei | 3 | 2337891 | A→G | R510R (CGT→CGC) | gyrA ← | 0.055 |
| cip | Δnth-Δnei | 3 | 2339173 | G→A | S83L (TCG→TTG) | gyrA ← | 1 |
| cip | Δnth-Δnei | 1 | 3164610 | C→T | R455H (CGT→CAT) | parC ← | 0.061 |
| cip | Δnth-Δnei | 3 | 3166096 | A→G | intergenic (-123/+15) | parC ← / ← ygiS | 0.082 |
| cip | Δnth-Δnei | 3 | 3166102 | A→C | intergenic (-129/+9) | parC ← / ← ygiS | 0.078 |
| cip | Δnth-Δnei | 2 | 986410 | A→C | S191S (TCT→TCG) | ompF ← | 0.055 |
| cip | Δnth-Δnei | 2 | 986411 | G→T | S191Y (TCT→TAT) | ompF ← | 0.055 |
| cip | ΔmutS | 1 | 2339173 | G→A | S83L (TCG→TTG) | gyrA ← | 1 |
| cip | ΔmutS | 2 | 2339173 | G→A | S83L (TCG→TTG) | gyrA ← | 1 |
| cip | ΔmutS | 3 | 2339173 | G→A | S83L (TCG→TTG) | gyrA ← | 1 |
| rif | ΔmutT | 1 | 4182782 | A→C | Q513P (CAG→CCG) | rpoB → | 1 |
| rif | ΔmutT | 2 | 4182782 | A→C | Q513P (CAG→CCG) | rpoB → | 1 |
| rif | ΔmutT | 3 | 4181140 | A→G | intergenic (+215/-105) | rplL → / → rpoB | 0.145 |
| rif | ΔmutT | 3 | 4182959 | T→G | I572S (ATC→AGC) | rpoB → | 1 |
| rif | ΔmutH | 1 | 4182790 | G→A | D516N (GAC→AAC) | rpoB → | 1 |
| rif | ΔmutH | 2 | 4182790 | G→A | D516N (GAC→AAC) | rpoB → | 1 |
| rif | ΔmutH | 3 | 4182791 | A→G | D516G (GAC→GGC) | rpoB → | 1 |

**Table S5:** *continued..*

| Antibiotic | Strains | Replicate | Position | Mutation | Annotation | Gene | Frequency |
|---|---|---|---|---|---|---|---|
| rif | ΔmutY | 1 | 4182779 | C→A | S512Y (TCT→TAT) | rpoB → | 1 |
| rif | ΔmutY | 2 | 4181680 | G→T | V146F (GTT→TTT) | rpoB → | 1 |
| rif | ΔmutY | 3 | 4181680 | G→T | V146F (GTT→TTT) | rpoB → | 0.843 |
| rif | ΔmutY | 3 | 4182779 | C→A | S512Y (TCT→TAT) | rpoB → | 0.075 |
| rif | ΔmutL | 1 | 4182790 | G→A | D516N (GAC→AAC) | rpoB → | 1 |
| rif | ΔmutL | 2 | 4182791 | A→G | D516G (GAC→GGC) | rpoB → | 1 |
| rif | ΔmutL | 3 | 4182790 | G→A | D516N (GAC→AAC) | rpoB → | 1 |
| rif | ΔmutL | 2 | 4185817 | T→C | R156R (CGT→CGC) | rpoC → | 0.069 |
| rif | ΔmutS | 1 | 4182778 | T→C | S512P (TCT→CCT) | rpoB → | 0.119 |
| rif | ΔmutS | 1 | 4182820 | C→T | H526Y (CAC→TAC) | rpoB → | 0.865 |
| rif | ΔmutS | 2 | 4182791 | A→G | D516G (GAC→GGC) | rpoB → | 1 |
| rif | ΔmutS | 3 | 4182778 | T→C | S512P (TCT→CCT) | rpoB → | 0.162 |
| rif | ΔmutS | 3 | 4182790 | G→A | D516N (GAC→AAC) | rpoB → | 0.849 |
| rif | ΔmutS | 3 | 4184958 | G→A | L1238L (CTG→CTA) | rpoB → | 0.067 |
| rif | ΔmutS | 3 | 4188547 | A→G | E1066E (GAA→GAG) | rpoC → | 0.098 |
| tet | ΔmutH | 1 | 1619399 | C→T | R94C (CGC→TGC) | marR → | 0.105 |
| tet | ΔmutH | 3 | 1619399 | C→T | R94C (CGC→TGC) | marR → | 0.236 |
| tet | Δnth-Δnei | 1 | 585602 | G→A | T11I (ACA→ATA) | ompT ← | 0.052 |
| tet | Δnth-Δnei | 3 | 585081 | T→A | N185Y (AAT→TAT) | ompT ← | 0.063 |
| tet | ΔmutL | 2 | 986173 | G→A | S270S (AGC→AGT) | ompF ← | 0.064 |
| tet | ΔmutL | 1 | 1619352 | T→C | L78P (CTG→CCG) | marR → | 0.172 |
| tet | ΔmutL | 2 | 1619352 | T→C | L78P (CTG→CCG) | marR → | 0.13 |
| tet | ΔmutS | 1 | 1619328 | C→T | A70V (GCA→GTA) | marR → | 0.676 |
| tet | ΔmutS | 2 | 1619234 | A→G | T39A (ACC→GCC) | marR → | 0.187 |
